# BCAT1 inhibition affects CD8^+^ T cell activation, exhaustion, and tumoral immunity by altering iron homeostasis

**DOI:** 10.1101/2023.02.25.530034

**Authors:** Francesca Lodi, Michelangelo Certo, Hagar Elkafrawy, Weixing Li, Hong A. Vu, Konstantin Gilbo, Li Su, Ian L. Pegg, Tobias Weiss, Marcel Bühler, Michael Weller, Charles Yeh, Jacob E. Corn, Kwon-Sik Park, Jeong-Hun Ko, Jacques Behmoaras, Claudio Mauro, Diether Lambrechts, Adonia E. Papathanassiu

## Abstract

The present study explores the role of the cytosolic branched chain amino acid aminotransferase (BCAT1) in CD8^+^ T cell activation, in general, and tumor immunity, in particular, and identifies a non-canonical function of the protein in iron homeostasis. Pharmacologic inhibition of BCAT1 using the novel drug ERG245 abrogates the effector functions of CD8^+^ T cells in vitro and metabolically reprograms the cells towards increased OXPHOS. In vivo, it suppresses activation of CD8^+^ T cells in DSS colitis leading to improved disease outcomes. Remarkably, withdrawal of BCAT1 inhibition further amplifies OXPHOS and gives rise to CD8^+^ T cells with increased cytotoxicity in vitro and in vivo. When combined with an anti-PD-1 treatment, temporal BCAT1 inhibition dramatically increases anti-PD-1 efficacy inducing complete and durable tumor regressions in the moderately immunogenic CT26 tumor model. Single cell RNA-seq data link expression of Bcat genes to exhausted T cells within the tumor microenvironment of human cancer patients, whereas in vitro assays indicate that BCAT1 inhibition partially prevents the adoption of a terminally exhausted phenotype by CD8^+^ T cells. We propose BCAT1 as a target for cancer combinatory therapies.

**SIGNIFICANCE:** The study explores for the first time the role of BCAT1 in CD8^+^ T cell activation and proposes novel strategies for using BCAT1 inhibitors in cancer and beyond. It demonstrates that BCAT1 exerts its function without significantly altering branched chain amino acid (BCAA) levels through a mechanism that controls iron homeostasis, a novel non-canonical mechanism of action, and implicates BCAT1 in the adoption of an exhausted phenotype by T cells found in human cancers. While the majority of metabolic drugs temper OXPHOS, it demonstrates that an agent that increases OXPHOS in CD8^+^ T cells can be used successfully as an immune-oncology drug.

## INTRODUCTION

Cancer immunotherapies, especially immune checkpoint inhibition (ICI) and CAR-T cell transfer, have become the new standard of care for several malignancies even though durable responses are observed only with a small subset of patients (1-4). Insufficient tumor immunogenicity, cancer immunoediting, irreversible T cell exhaustion, and a highly immunosuppressive tumor microenvironment (TME) are considered some of the mechanisms contributing to primary and acquired resistance to ICI and CAR-T therapies (2,4). It is well understood that tumor-infiltrating immune cells encounter a hostile tumor microenvironment (TME), characterized by areas of low pH and hypoxia, a modified extracellular matrix that impedes immune cell migration, the presence of immunosuppressive cytokines such as TGF-β, and the lack of nutrients due to dysregulated cancer cell metabolism (5). Adoptive cell transfer (ACT) of cancer specific CD8^+^ T cells in syngeneic tumor models has indicated that T cells rapidly become dysregulated upon entering the TME (6), whereas advances in single cell (sc) technologies have provided evidence that dysfunctional CD8^+^ T cells found in the TME are transcriptionally and functionally diverse, reside along a spectrum of dysfunction, and follow predictable trajectories of transformation from a naïve to a terminally exhausted phenotype (7-10). Studies integrating therapeutic responses in patients with sc transcriptomic analysis of tumoral T cells from relevant samples have given rise to a consensus that terminal exhausted CD8^+^ T cells, which are typically refractory to anti-PD-1 (αPD1) treatment, exhibit specific characteristics such as: a) the presence of high levels of PD-1, b) the co-expression of additional inhibitory molecules (e.g., TIM3 and LAG3) and cytotoxic molecules such as various granzymes, and c) the ability to constitutively secrete CXC ligand 13 (CXCL13) (9,10). Not surprisingly, several studies have suggested that distinct functional states of CD8^+^ T cells found within the TME are associated with distinct metabolic programs (10,11), although it is unclear if metabolic drivers exist that propel the cells along the dysfunctionality axis.

The classical understanding of the role of metabolism in T cell activation centers on the idea that adoption of a specific phenotype by the cells requires some form of metabolic reprogramming. Naïve cells have limited energy needs that are primarily met through mitochondrial oxidative phosphorylation (OXPHOS). T-cell receptor (TCR) activation initiates aerobic glycolysis, which is subsequently maintained through CD28 co-stimulation. Aerobic glycolysis is the preferred energy source of effector T cells. It is regulated by the PI3K-AKT-mTORC1 and Myc signaling pathways (13-15). Transition of CD8^+^ T cells from effector to memory cells is linked to the reactivation of OXPHOS as the primary route of ATP generation (16,17). CD8^+^ T cells expressing PD-1 generally exhibit enhanced fatty acid oxidation of endogenous lipids, suppressed glycolysis and impaired OXPHOS (11). Moreover, inhibition of OXPHOS by electron transport chain (ETC) inhibitors in activated T cells has been shown to suppress the self-renewal capacity of the cells and to upregulate biomarkers of exhaustion (18). More recently, studies have suggested that glycolysis and OXPHOS are not mutually exclusive and that a reciprocal and well-coordinated relationship exists between these two processes. For example, a significant contribution of OXPHOS in ATP production was reported for effector T cells grown ex vivo but not for effector cells generated in vitro (19). Yet, in vitro, at the very early stages of activation prior to clonal expansion, CD8^+^ T cells exhibit peak metabolic rate with enhanced glycolysis and OXPHOS (20,21). A similar phenomenon has been reported for a CD8^+^ T cell population, which exhibited biomarkers of terminal exhaustion, was found present in the tumors and peripheral blood of melanoma patients, and was associated with resistance to ICI (10). In that population, glycolysis and OXPHOS were both elevated although the increase in OXPHOS was preeminent and correlated with the rise of a terminal exhaustion phenotype. A relationship between the metabolic profile of exhausted T cells and their spatial organization appears to exist with highly metabolic PD-1^+^-expressing CD8^+^ T cells found near tumors and moderately metabolic ones relegated to the periphery (11). Collectively, the above studies suggest that tumoral CD8^+^ T cell function is moderated by the interplay of distinct metabolic pathways thus raising the possibility of overcoming ICI resistance by identifying relevant druggable metabolic drivers.

Metabolism of branched chain amino acids (BCAA) such as leucine (Leu) is initiated by BCAT isozymes, which catalyze the reversible transfer of an amino group from BCAAs to α-ketoglutarate (αKG) to generate glutamate and the relevant α-keto acids. In the past decade, numerous studies have identified a role for the cytosolic isoform (BCAT1) in the detection, prognosis, progression, and development of multi-drug resistance of different types of cancer but have failed to specify a unified mechanism of action (22-25). BCAT1-driven restriction of αKG leading to altered ten-eleven translocation methylcytosine dioxygenase (TET) activity and DNA hypermethylation, mTORC1 regulation, autophagosome synthesis, and activation of cancer type-specific oncogenes have been proposed (22-25). These mechanisms have been considered downstream of BCAA catabolism or BCAA synthesis. More recently, evidence have emerged indicating that BCAT1 and its mitochondrial counterpart BCAT2 potentially possess non-canonical functions (26-28). These functions are thought to be mediated through certain structural elements, unique to BCATs among other aminotransferases; these elements include a redox active CXXC motif, present near the active site of the enzymes, and the existence of putative phosphorylation, acetylation, and palmitoylation sites (26,29,30). It has been proposed that the cellular behavior of BCAT1 is regulated by the redox environment and post-translational modifications, which control translocation of the protein to distinct cellular compartments (including the membrane and the nucleus) as well as complexing with different binding partners (26). Despite the advances in understanding the role of BCAT1 in cancer in general, its role in immuno-oncology and especially in CD8^+^ T cell biology remains unexplored.

Here, we report that: a) BCAT1 is expressed by CD8^+^ T cells, b) inhibition of the protein or gene deletion abolishes effector functions during activation of naïve CD8^+^ T cells, and c) inhibition of effector functions does not correlate with BCAA metabolism. Newly activated CD8^+^ T cells that have experienced BCAT1 inhibition (BCAT1i) exhibit dramatically increased levels of OXPHOS, once BCAT1i is withdrawn. The metabolic reprogramming of the cells with temporal BCAT1i ultimately yields CD8^+^ T cells with increased effector functions in vitro and in vivo. Furthermore, we show that BCAT1i dramatically synergizes with αPD1 therapy to yield complete tumor regressions in a moderately immunogenic murine tumor model. Using sc-RNAseq data obtained from human tumor specimens, we demonstrate that genes of the BCAT isozymes are preferentially expressed by terminally exhausted T cells and that BCAT1i partially suppresses the development of CD8^+^ T cell exhaustion in vitro. Mechanistically, we identify a rapid increase in the levels of mitochondrial iron following BCAT1i and speculate that BCAT1 partners with iron regulatory protein 1 (IPR1) to regulate iron homeostasis at the subcellular level.

## RESULTS

### BCAT1 is required for differentiation of naïve CD8^+^ T cells to effector T cells

Naïve CD8^+^ T cells express BCAT1 at genomic and proteomic levels that remain unaltered upon activation with anti-CD3 and anti-CD28 antibodies (αCD3/αCD28) (Figs. 1A, S1A, and S1B). Human CD8^+^ (hCD8^+^) T cells do not show a preference for Bcat1 expression and both Bcat isoforms are detected at similar genomic levels in naïve and activated cells by qPCR (Fig S1B).

**Figure 1.**
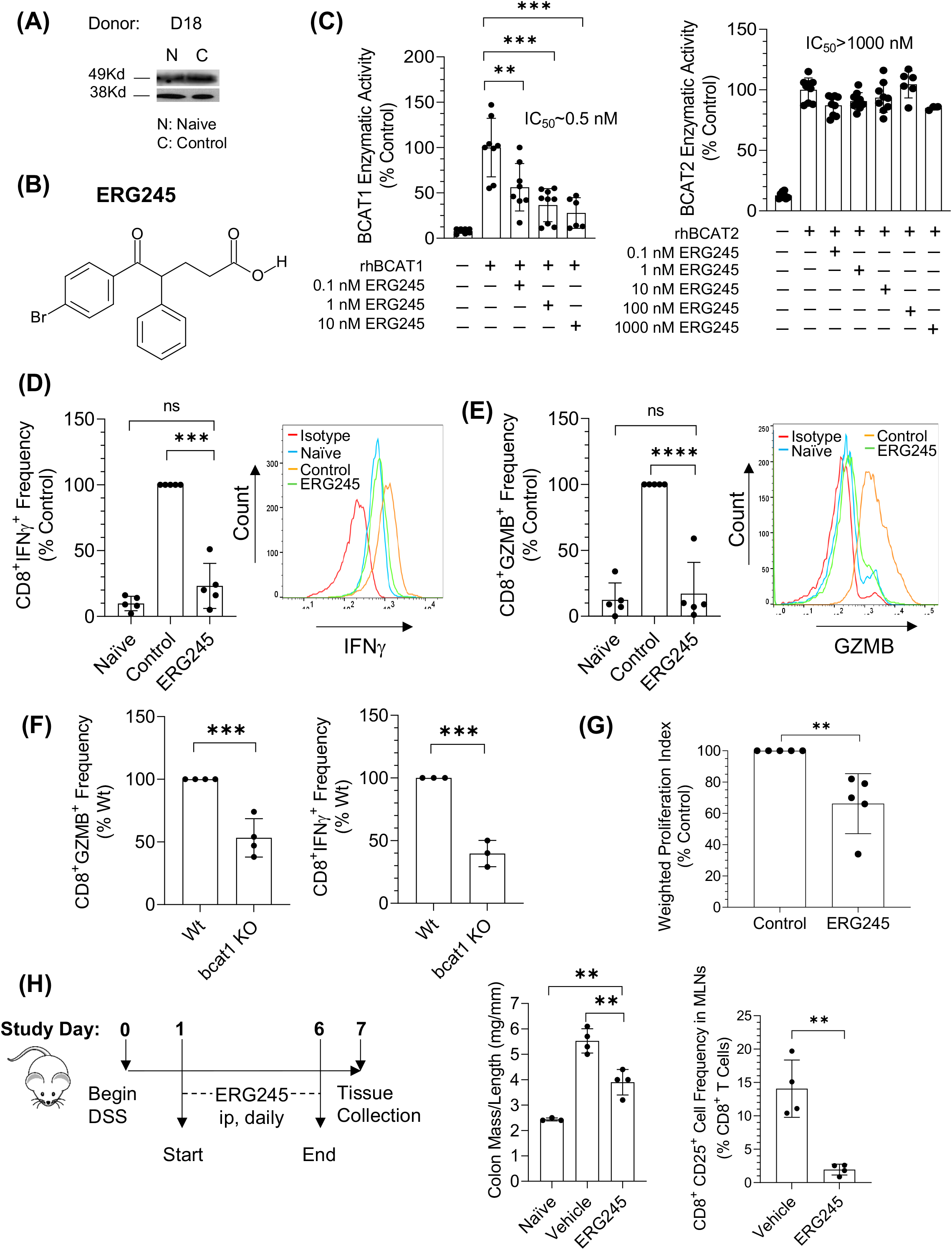
(A) BCAT1 protein expression in naïve and control (activated with αCD3/αCD28 antibodies) hCD8^+^ T cells. Three individual donors. (B) The structure of BCAT1 inhibitor ERG245. (C) Inhibition of rhBCAT1 (left) and rhBCAT2 (right) enzymatic activities by ERG245. Two independent experiments performed in triplicate. (D) IFNγ expression following activation of naïve hCD8^+^ T cells in the presence and absence of ERG245. Average of five individual donors (Left). Example of a flow cytometry analysis of a single donor (Right). (E) GZMB expression following activation of naïve hCD8^+^ T cells in the presence and absence of ERG245. Average of five individual donors (Left). Flow cytometry analysis of a single donor (Right). (F) GZMB and IFNγ expression after activation of mCD8^+^ T cells isolated from the spleens of age-matched wt and bcat1 KO mice. Three or four individual pairs. (G) Proliferation of activated hCD8^+^ T cells treated with ERG245. Weighted Proliferation Index is defined in Materials and Methods. (H) Experimental design, disease outcome, and frequency of newly activated CD8^+^CD25^+^ T cells present in the MLNs of C57/BL6 mice experiencing DSS-colitis and treated with either Vehicle control or 5 mg/kg ERG245.

To explore the role of BCAT1 in CD8^+^ T cells, we synthesized a series of BCAT1 inhibitors and selected the novel compound ERG245 (Fig. 1B) for further study. ERG245 inhibited the enzymatic activity of recombinant human BCAT1 (rhBCAT1) with an IC_50_ value of ∼0.5 nM and showed >2000-fold selectivity for BCAT1 over BCAT2. (Fig. 1C). ERG245 also exhibited acceptable pharmacokinetic properties, suitable for use of the drug in in vivo experiments (Table S1). We then subjected naïve hCD8^+^ T cells, isolated from the peripheral blood of 5 individual donors, to αCD3/αCD28 activation with and without BCAT1i using ERG245 and analyzed the expression of effector biomarkers using flow cytometry. In Figs 1D and 1E, exposure of the cells to ERG245 completely suppressed production of IFNγ and granzyme B (GZMB) in activated CD8^+^ T cells. The role of BCAT1 in CD8^+^ T cell differentiation was corroborated using murine CD8^+^ (mCD8^+^), isolated from the spleens of age-matched wt and bcat1 KO mice (Fig. 1F). In addition, BCAT1i was associated with partial inhibition of CD8^+^ T cell expansion (Fig. 1G). To determine if BCAT1 exhibits a similar role in vivo, we examined the effect of ERG245 on the disease activity and activation status of CD8^+^ T cells, found in mesenteric lymph nodes (MLNs) of mice suffering from acute dextran sulfate sodium (DSS)-induced colitis, a model of inflammatory bowel disease (IBD) (31). It has been reported that, during colitis and other gut diseases, the presence of bacteria in intestinal and colonic tissues induces migration of dendritic cells (DCs) to MLNs, and subsequent priming of T cells (32). In the experiment of Fig. 1H, colitis was induced in mice through the continuous consumption of drinking water containing DSS and disease activity was assessed by determining the ratio of colon mass/colon length at the end of the experiment. That ratio typically increases following DSS administration as a result of colonic-inflammation and inflammation-driven fibrosis. In Fig. 1H, daily administration of ERG245 (starting 24h after disease induction) mitigated disease activity and decreased the frequency of CD8^+^CD25^+^ T cells present in the MLNs. Induction of CD25, also known as interleukin-2Rα, is an early sign of CD8^+^ T cell activation in vitro and in vivo (33).

### Temporal BCAT1i yields CD8^+^ T Cells with increased cytotoxicity

To determine if the effect of BCAT1i on CD8^+^ T cell effector functions was reversible, the cells were exposed to ERG245 for 24h at the beginning of activation. Subsequently, conditioned media (CM) were replenished with fresh media containing IL-2 and cells were cultured for an additional 72h in the absence of BCAT1i. At the end of the experiment, the cell surface expression of the cytotoxic biomarker GZMB was determined by flow cytometry. Fig. 2A indicates that, not only the effects of BCAT1i on effector functions of CD8^+^ T cells were reversible, but also withdrawal of ERG245 led to the generation of T_eff_ cells with increased cytotoxicity. That observation suggested that it might be possible to exploit temporal BCAT1i in the context of cancer in order to increase the cytotoxicity of CD8^+^ T cells that are being re-invigorated by αPD1 therapies. To explore the idea, we employed the murine CT26 syngeneic colon cancer model. CT26 tumors are moderately immunogenic and αPD1 therapies have only a modest or no effect on tumor growth and survival in the model. In our hands, the effect of αPD1 therapy on CT26 tumor growth was dependent on the tumor volume at the time of treatment initiation with early applications of checkpoint inhibition being more effective than later ones. This was in line with published reports describing a dynamic CT26 tumor microenvironment (TME) possessing an immune cell infiltrate that varied over time and was responsive to αPD1 therapy only when animals were dosed within a certain timeframe (34,35). In the CT26 model, temporal BCAT1i was employed in vivo in combination with αPD1 therapy as shown in the experimental design of Fig. 2B. The treatment regimen was initiated when the tumor volumes were ∼50 mm^3^. In Fig. 2B, brief treatment of CT26-tumor bearing animals with ERG245 alone did not alter the kinetics of tumor growth nor did it improve survival. On the other hand, αPD1 monotherapy had a limited therapeutic effect with only 1/6 animals experiencing complete tumor regression (CTR). In contrast the combination treatment had a dramatic impact with 5/6 animals experiencing CTR (Figs. 2B and 2C). Animals exhibiting CTR at day 49 of treatment were re-inoculated with CT26 tumor cells on the opposite flank (Fig. 2B). No evidence of tumor presence was observed when the experiment was terminated on day 63 of treatment suggesting the possibility that immunological memory was established.

**Figure 2.**
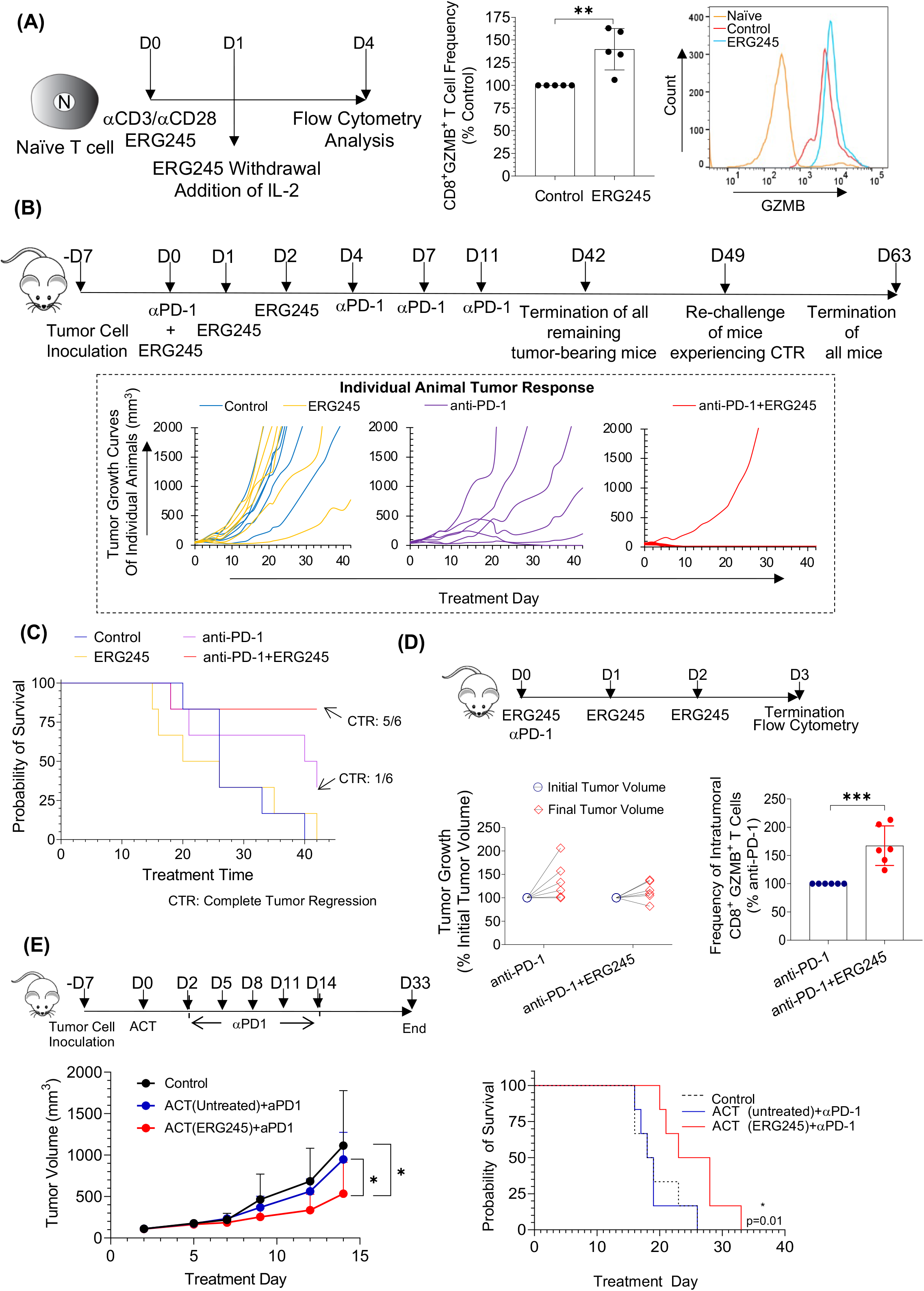
(A) Cytotoxicity of hCD8^+^ T cells following a brief (24h) BCAT1i by ERG245 at the beginning of activation, withdrawal of the drug, and additional culturing of the cells for 72h in the presence of media containing IL-2. From left to right: depiction of the experimental design, frequency of CD8^+^GZMB^+^ T cells at 96h (5 individual donors), and an example of flow cytometry analysis at 96h (single donor). (B) In vivo experiment comparing the efficacy of the ERG245+ αPD1 combination to that of αPD1 monotherapy. From top to bottom, depiction of the experimental design and individual tumor response data. Four groups (n=6 animals/group): Control (control ab; same regimen as αPD1), ERG245 (5 mg/kg, ip, bid, given on days 0, 1, and 2 of treatment), anti-PD-1 (10 mg/kg αPD1, ip, qd, given on days 0, 4, 7, and 11 of treatment), and ERG245+anti-PD-1. Complete tumor responses (CTR) were observed in the anti-PD-1 group (1/6 or 17%) and in the ERG245+anti-PD-1 (5/6 or 83%). (C) Kaplan-Meier survival curve associated with the experiment described in Fig. 2B. (D) In vivo experiment examining the presence of cytotoxic (CD8^+^GZMB^+^) T cells in the TME of animals treated with either anti-PD-1 monotherapy (10 mg/kg αPD1, ip, a single dose on day 0 of treatment) or a combination of ERG245 (5 mg/kg, ip, bid, given on days 0, 1, and 2 of treatment) and anti-PD-1. From top to bottom and left to right, depiction of the experimental design, individual tumor volumes at the beginning and end of the test, and frequency of CD8^+^GZMB^+^ T cells present in the TME of the animals. Two independent experiments, 6 individual animals per condition matched according to initial tumor volumes. (E) Efficacy of ACT of CD8^+^ T cells (untreated or ERG245-treated for 24h) combined with αPD1 therapy. From top to bottom and left to right, depiction of the experimental design, tumor growth prior to first observed death on day 16 of treatment, and Kaplan-Meier survival curves associated with the experiment. Control group: untreated animals.

To verify that BCATi increases the cytotoxicity of tumor infiltrating CD8^+^ T cells that are being re-invigorated by αPD1 therapy, we immunophenotyped the TMEs of CT26 tumors 96h after they received a single dose of αPD1 with or without the concomitant administration of ERG245 for 3 days. The experimental design and the combined data from two individual experiments are shown in Fig. 2D. Addition of ERG245 to the αPD1 treatment increased the frequency of CD8^+^GZMB^+^ T cells in the CT26 TME by >60%. To ensure that the effect was independent of the initial tumor volume (and thus the specific composition of CT26 TMEs), the experiments included tumors ranging from ∼100 to ∼800 mm^3^ (Fig. 2D). Addition of ERG245 to αPD1 consistently increased the frequency of cytotoxic cells present in the CT26 TME across all tumor volumes tested. To ascertain that the increase in CD8^+^GZMB^+^ T cell frequency was due to a direct effect of ERG245 on CD8^+^ T cells, we combined αPD-1 therapy with adoptive cell transfer (ACT) of splenic CD8^+^ T cells, previously activated with αCD3/αCD28 for 24h in the presence or absence of ERG245. The treatment was initiated when the tumor volumes of the mice was ∼100 mm^3^. In Fig. 2E, the ACT of Vehicle-treated CD8^+^ T cells combined with αPD1 therapy exhibited identical tumor growth kinetics and probability of survival of CT26-tumor bearing animals to the ones observed for control (untreated) animals. In contrast, the ACT of ERG245-treated CD8^+^ T cells along with αPD1 therapy decreased tumor growth and increased the median survival of animals by ∼40% (Fig. 2E).

### BCAT1i results in irreversible upregulation of OXPHOS

To understand the mechanism of action of BCAT1 in CD8^+^ T cells, we first interrogated the metabolic effect of BCAT1i or gene deletion in hCD8^+^, mCD8^+^, and Jurkat cells, activated in SILAC media containing ^15^N-Leu. Cells with fully functional BCAT1, operating in the forward action (see Supplementary Fig. S1C), are expected to transfer a ^15^N-labeled amino group from a labeled Leu molecule present in the cytosol to an unlabeled Glu molecule. In the experiments of Fig. 3A, ^15^N-Glu was detected only in the cell lysates of untreated Jurkat cells but not in those of untreated hCD8^+^ or wt mCD8^+^ cells. As expected, exposure of wt Jurkat cells to ERG245 or deletion of Bcat1 gene using the CRISP-Cas9 system resulted in reduced levels of ^15^N-Glu compared to control cells, demonstrating the forward enzymatic action of BCAT1 in those cells (Fig. 3A). In hCD8^+^ as well as mCD8^+^ T cells, ^15^N-Leu accounted for >75% of intracellular Leu at 24h of activation indicating that the cells were able to readily uptake Leu from their extracellular environment but did not utilize it to generate Glu. Interestingly, total intracellular levels of Leu (^14^N- and ^15^N-Leu) as well as isoleucine (Ile) increased significantly in hCD8^+^ T cells exposed to ERG245 but not in mCD8^+^ from bcat1 KO animals (Fig. 3B)

**Figure 3.**
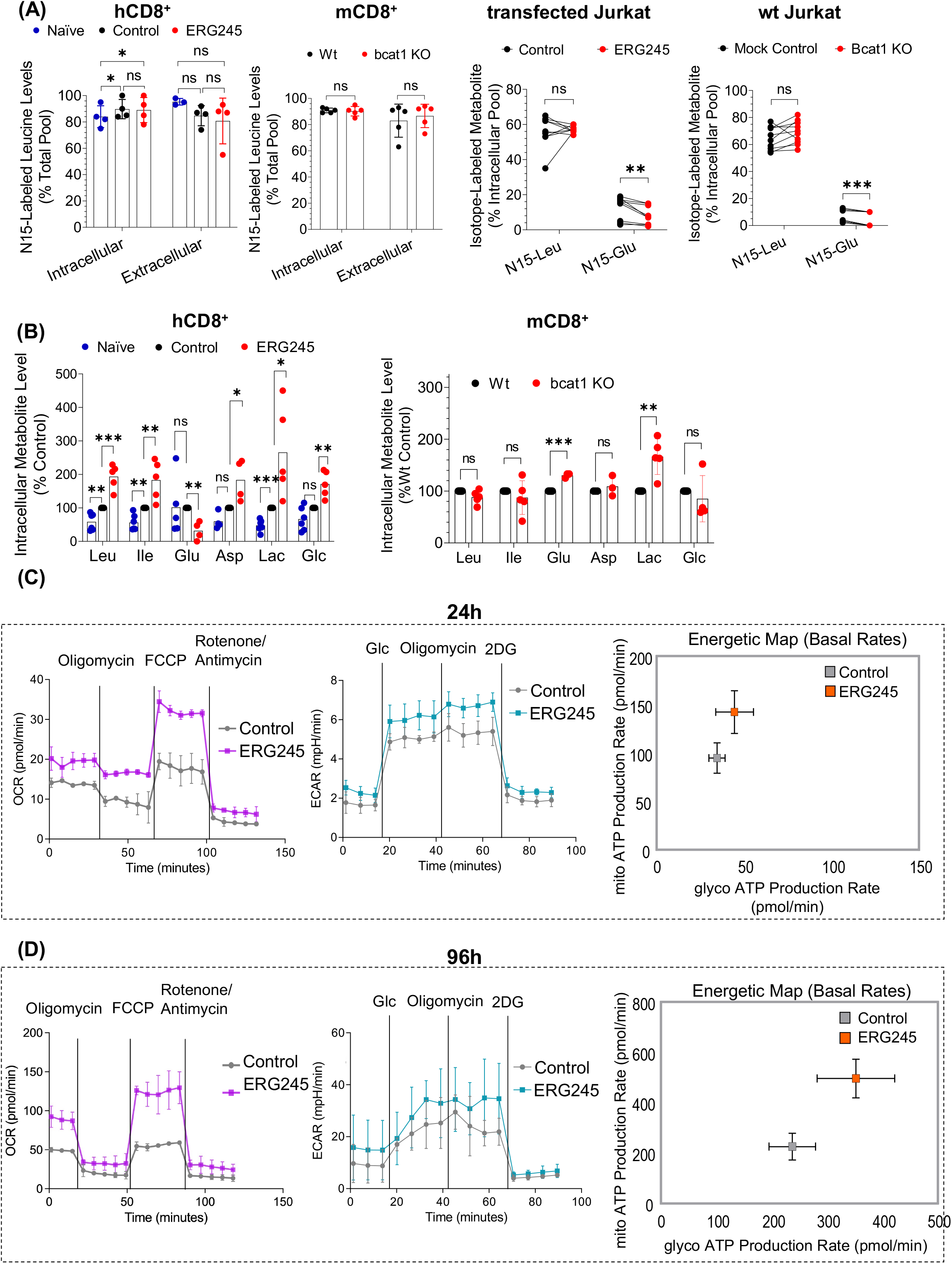
(A) Intracellular or extracellular levels of ^15^N-Leu present in hCD8^+^ (4 individual donors), mCD8^+^ (5 individual age-matched wt and bcat1 KO animals), and Jurkat cells (two individual experiments performed in triplicate or quadruplicate) cultured in Silac media containing 50 mg/L ^15^N-Leu and activated with αCD3/αCD28 for 24h (CD8+ T cells) or 3h (Jurkat cells) prior to analysis. Levels of ^15^N-Leu are depicted as % Total Pool (^14^N- and ^15^N-Leu). Intracellular levels of ^15^N-Glu are also shown for Jurkat cells. No ^15^N-Glu was detected at 24h for hCD8^+^ or mCD8^+^ T cells by GC-MS. (B) Total intracellular levels of various metabolites in the hCD8^+^ and mCD8^+^ experiments. (C) Seahorse analysis of αCD3/αCD28-activated hCD8^+^ T cells treated with ERG245 for 24h. Four individual donors performed in triplicate. From left to right: Oxygen consumption rate (OCR), Extracellular acidification rate (ECAR), and depiction of combined production of mitochondrial ATP and glycolytic ATP. (D) Seahorse analysis of αCD3/αCD28-activated hCD8^+^ T cells treated with ERG245 as described in Fig. 2A: Four individual donors performed in triplicate. From left to right: Oxygen consumption rate (OCR), Extracellular acidification rate (ECAR), and depiction of combined production of mitochondrial ATP and glycolytic ATP.

Intracellular Leu has a well-documented role in mTOR activation and mTOR activation is linked to T cell proliferation, effector cell differentiation, and changes in cell metabolism (36). Subsequently, we examined the effect of BCAT1i on mTOR activation. Supplemental Fig. S2D confirms that activation of hCD8^+^ T cells with αCD3/αCD28 antibodies leads to activation of mTOR, as determined by an increase in the phosphorylation levels of mTOR and its downstream target ribosomal protein S6. Activation of mTOR was not affected by exposure of the cells to ERG245 as shown in Fig. S2D. Similarly, ERG245 treatment did not affect the levels of cMYC found in activated hCD8^+^ T cells (Fig. S1E). c-MYC is a transcription factor required for the metabolic reprogramming of activated T lymphocytes (37). Analysis of the intracellular (Fig. 3B) and extracellular (Fig. S1F) levels of lactate (Lac) and glucose (Glc) in both ERG245-treated hCD8^+^ and mCD8^+^ Bcat1 KO T cells further cooroborated that the cells successfully underwent metabolic reprogramming towards aerobic glycolysis following αCD3/αCD28 activation despite inhibition or deletion of BCAT1. Details of this analysis can be found in the Supplementary Text.

We then sought to examine the effect of BCAT1i on CD8^+^ T cell bioenergetics. We performed Seahorse analysis of hCD8^+^ T cells, activated in the presence or absence of ERG245 for 24h (Fig. 3C). BCAT1i led to an increase in oxygen consumption rate (OCR) of activated hCD8^+^ T cells without an effect on the extracellular acidification rate (ECAR) of the cells. As a result, ERG245-treated CD8^+^ T cells exhibited an apparent trend of increased ATP production in the mitochondria via OXPHOS (Fig. 3C). To explore the reversibility of the trend upon BCAT1i withdrawal, we analyzed the bioenergetics status of the cells 72h after removal of ERG245 treatment. Fig. 3D indicates that, not only the increase in OXPHOS was irreversible, but it was also amplified upon withdrawal of BCATi. The rate of ATP production via OXPHOS increased by 2-fold in hCD8^+^ T cells treated briefly with ERG245 compared to untreated control and the increase was statistically significant, whereas the rate of glycolytic ATP production remained essentially unaltered. Additional data from the Seahorse analysis at 24h and 96h illustrate that BCAT1i increased the basal and maximal respiration of the CD8^+^ T cells without affecting their spare respiratory capacity (Figs. S1I and S1J).

### BCAT1 regulates mt iron levels upstream from OXPHOS increase

To further explore the mechanism of action of BCAT1 in hCD8^+^ T cell activation, we determined the transcriptomic changes induced by ERG245 treatment at 24h (continuous inhibition) and 96h (temporal inhibition). Using bulk RNA-sequencing (RNA-seq), we identified 837 genes that were differentially expressed (DEGs) after BCAT1i for 24h (Figs. 4A and S2A). In the Volcano plot of Fig. 4A, the genes mostly affected by ERG245 treatment belong to a family of mt encoded genes, MTRNR2, related to humanin. Humanin is a mt peptide, known to protect cells from oxidative stress and to increase mitochondrial respiration rate and biogenesis (38). DEGs that were either upregulated (Fig. 4B) or downregulated (Fig. 4C) were visualized as protein-protein interaction (PPI) networks using Cytoscape stringAPP. Figure 4B suggests that BCAT1i triggers a DNA damage response along with pro- and anti-apoptotic pathways. An increase in the expression of genes involved in [Fe-S] cluster biogenesis is also observed. Significantly downregulated genes were further clustered using the MCODE algorithm (Fig. 4C). The clustering revealed that BCAT1i affected primarily cell cycling genes. Processes associated with protein translation such as mRNA splicing and ribosome biogenesis were also downregulated along with genes linked to immune response and, cholesterol biosynthesis, and glycolysis. Pathway enrichment analyses were also performed using the KEGG (Fig. 4D) and Reactome databases (Fig. S2B). Both databases indicated an enrichment in pathways associated with cell cycling and DNA replication with KEGG additionally suggesting that DNA repair pathways (such as Fanconi anemia, homologous recombination, and pyrimidine biosynthesis) were affected.

**Figure 4.**
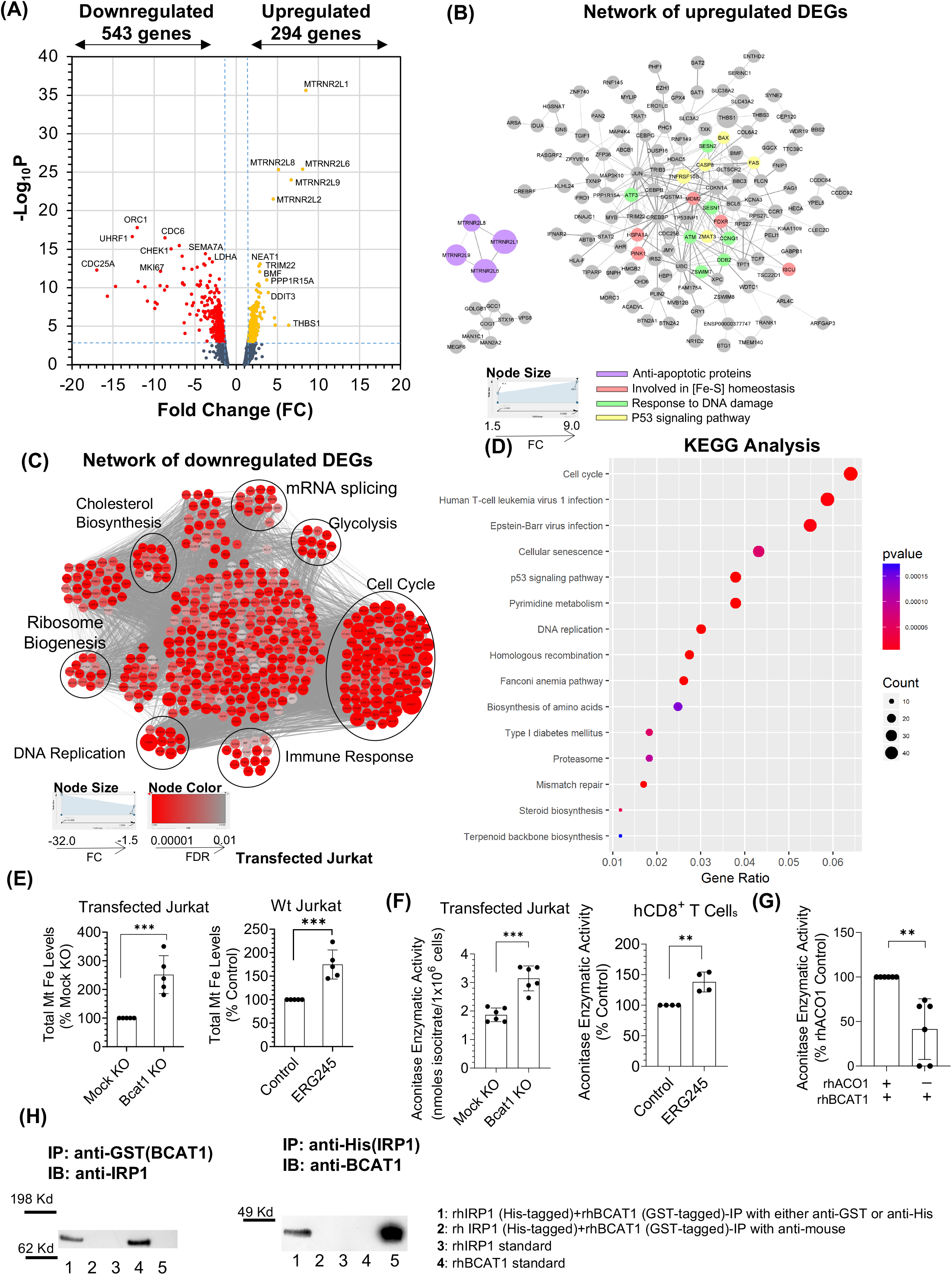
(A) Volcano plot of genes present in hCD8^+^ T cells from 3 individual donors activated in the presence and absence for ERG245 for 24h. DEGs defined as: FPKM >10, |FC|≥1.5 and p<0.001. (B) Visualization of a PPI network among upregulated DEGs with FDR<0.01. The network was created using the Cytoscape stringAPP. Highlighted genes are associated with the annotated biological processes. The node size varies with FC as shown. (C) Visualization of a PPI network among downregulated DEGs with FDR<0.01. The network was created using the Cytoscape stringAPP and clustered using the MCODE clustering algorithm. Color intensity of the nodes varies with FDR as shown. The size of the nodes varies with FC as shown. (D) A dotplot showing the top 15 KEGG pathways, found to be significantly altered after ERG245 treatment. (E) Iron levels in the mt of transfected (left) or wt (right) Jurkat cells following activation for 60 min. (F) Levels of cytosolic aconitase enzymatic activity in transfected Jurkat cells or hCD8^+^ T cells treated with ERG245. (G) Aconitase activity of rhACO1 measured in mixtures without and with rhBCAT1 and in the presence of activating solutions containing (NH_4_)Fe(SO_4_)_2_ and cysteine. (H) Immunoprecipitation experiments involving mixture of rhIRP1 tagged with His and rhBCAT1 tagged with GST and performed in the absence of an activating solution containing (NH_4_)Fe(SO_4_)_2_ and cysteine as described in Materials and Methods. IP: immunoprecipitation. IB: immunoblotting.

To further probe the mechanism of action of BCAT1i, we performed a weighted gene co-expression network analysis (WGCNA) on the entire RNA-seq gene dataset and identified possible hub genes controlling specific functional modules (see Bioinformatics section in Supplementary Text). We have highlighted two such hub genes: Cisd1, whose loss after ERG245 treatment was linked to suppression of ribosome biogenesis by WGCNA and Gene Ontology (GO), and Iscu, whose increase after ERG245 treatment was linked to induction of ER stress (Tables S3 and S4, respectively). Interestingly, both genes are located in the mt and are involved in [Fe-S] cluster biogenesis. CISD1 (or mitoNEET) is located on the outer mt membrane and is known to transfer its [2Fe-2S] clusters to cytosolic proteins such as IRP1, whereas ISCU acts as a scaffold for the assembly of [2Fe-2S] clusters in the mt (39,40). The effect of ERG245 on these genes raises the possibility that BCAT1i alters mt iron homeostasis. To explore the above hypothesis, we measured iron concentrations in the mitochondria of wt Jurkat cells in the absence and presence of ERG245, 60 min after activation with αCD3/αCD28 antibodies. We performed similar measurements with Jurkat Mock KO and Jurkat Bcat1 KO cells. As shown in Fig. 4E, BCAT1i led to a rapid increase in mt iron and that phenotype was reproducible with Bcat1 gene deletion. Because the transfer of [2Fe-2S] clusters from CISD1 to IRP1 converts the latter into [4Fe-4S] cluster-containing aconitase 1 (ACO1), we examined the effect of BCAT1i on the cytosolic aconitase activity of the cells. In Fig. 4F, loss of BCAT1 function led to an increase in ACO1 activity. Considering that aconitase enzymatic assays require chemical activation with a solution containing (NH_4_)Fe(SO_4_)_2_ and cysteine in order to mitigate for the oxidation of ACO1 [4Fe-4S] clusters during cell lysis, we examined the possibility that the presence of rhBCAT1 interferes with the chemical reconstitution of the [4Fe-4S] clusters in apo-IRP1. To test that, we measured the aconitase activity of rhIRP1 protein, chemically activated in the absence or presence of rhBCAT1. The results of Fig. 4G suggest that rhBCAT1 limits chemical activation of apo-IRG1, possibly by forming a physical complex with the protein. To examine the latter, we incubated rhIRP1 and rhBCAT1 under non-activating conditions and immunoprecipitated either IRP1 or BCAT1 with an appropriate antibody. Fig. 4H suggests that BCAT1 is able to physically interact with apo-IRP1. In contrast, under conditions of chemical activation, rhACO1 did not complex with rhBCAT1, although it was able to complex with rhBCAT1 that was inhibited with ERG245 (Supplementary Fig. S2G).

Transcriptomic analysis using bulk RNA-seq was also performed after temporal exposure of CD8^+^ T cells to ERG245 (Fig. 5A and Fig. S3A). The volcano plot of Fig. 5A identified 986 genes that were differentially expressed after temporal BCAT1i. GSEA analysis, performed on the two RNA-seq genesets in their entirety, revealed that ERG245 withdrawal reversed aspects of the phenotype observed after 24h exposure to the drug, especially the expression of genes targeted by Myc as well as the genes involved in the process of OXPHOS (Fig. S3B). Overall, 71 genes were commonly found in the DEGs of both groups (Fig. 5C). From these genes, 49 genes reversed and 22 genes maintained expression direction (Fig. 5C and Supplementary Table S5). Most notably, Bcat1 remained downregulated in both groups as did the proliferation marker Mki67, suggesting a lasting cell growth inhibition despite the increased cytotoxicity observed in Fig. 2B. DEGs that were upregulated following temporal exposure to ERG245 formed a PPI network comprising clusters of genes primarily linked to ribosome biogenesis and OXPHOS (Fig. 5D). The network of Figure 5D also included clusters of genes associated with NK cytotoxicity and glutathione metabolism. A similar analysis of DEGs that were downregulated in the treated cells revealed only a single gene cluster associated with histone modification (Fig. 5E).

**Figure 5.**
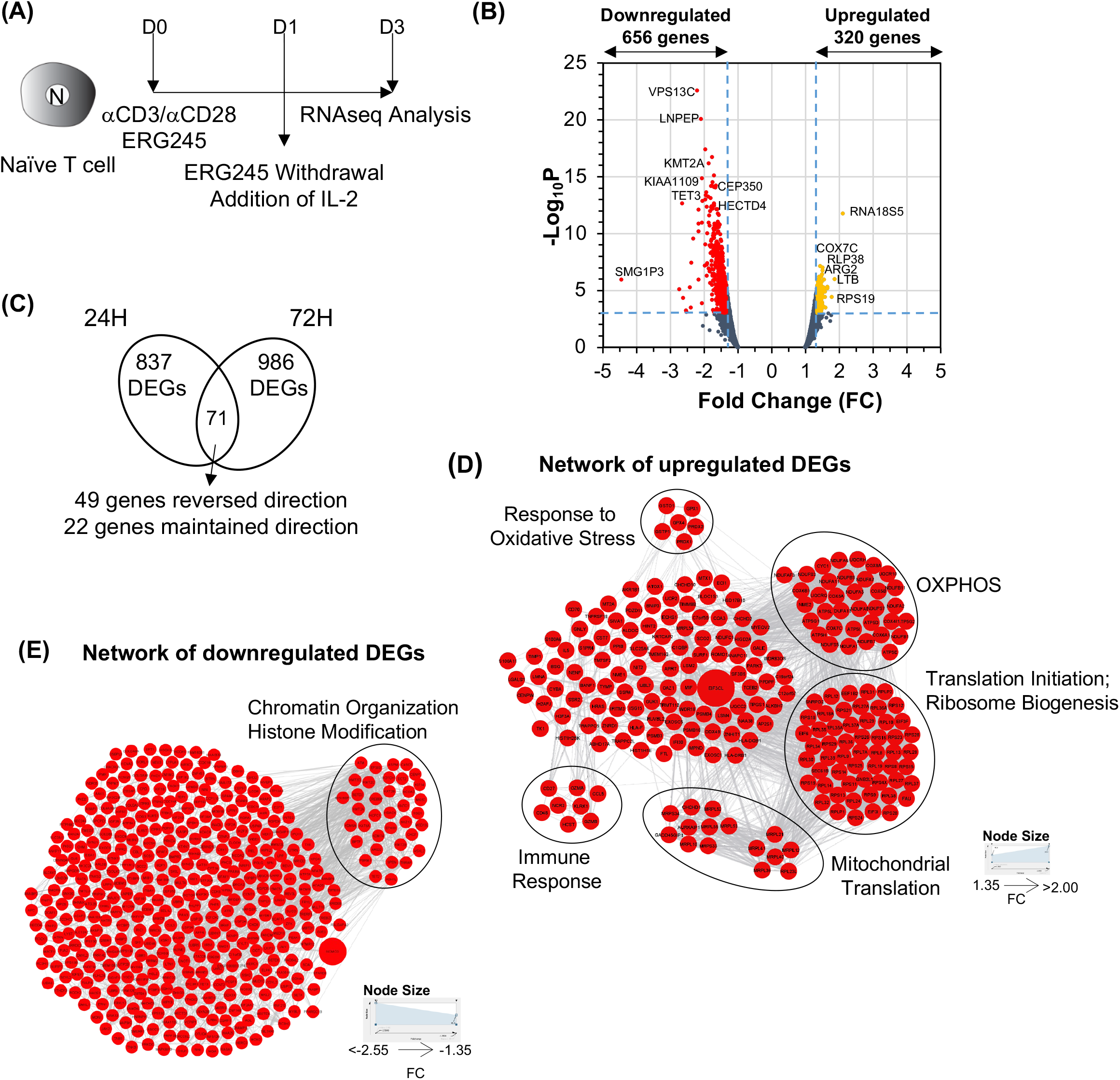
(A) Experimental design of the RNAseq experiment at 72h. (B) Volcano plot of genes present in αCD3/αCD28-activated hCD8^+^ T cells from 3 individual donors treated with and without ERG245, as depicted in Fig. 2A. DEGs defines as: FPKM >10, |FC|≥1.35 and p<0.001. (C) Venn diagram illustrating common genes in the DEGs of the RNAseq data at 24h and 72h (D) Visualization of a PPI network among upregulated DEGs with FDR<0.01. The network was created using the Cytoscape stringAPP and clustered using the MCODE clustering algorithm. The size of the nodes varies with FC as shown. (E) Visualization of a PPI network among downregulated DEGs with FDR<0.01. The network was created using the Cytoscape stringAPP and clustered using the MCODE clustering algorithm. The size of the nodes varies with FC as shown.

### BCAT1i Reverses Terminal Exhaustion in CD8^+^ T cells

To evaluate BCAT1 as a druggable target in human cancers generally and in immune-oncology specifically, we examined the expression of both Bcat isoforms in the scRNA-seq dataset of Ref 41. In that study, Qian et al. analyzed normal and tumor tissue samples from 36 treatment-naïve patients with 4 different types of cancer, specifically colon cancer (CRC, n=7), ovarian cancer (OvC, n=5), breast cancer (BC, n=16), and lung cancer (LC, n=8), and clustered individual cells per cell and tissue type. We first interrogated cells belonging to an individual cancer type at the level of the various cell clusters. The results are shown in Figs. 6A, 6B and S4A. Collectively, the data suggested that Bcat1 was mainly expressed by myeloid cells. These results were in agreement with previous observations reporting preferential expression of BCAT1 in primary human macrophages (42). Among the myeloid cells from CRC (n=3,660), OvC (n=6,526), BC (n=1,965), and LC (n=22,535), Bcat1 expression was detected in approximately 28%, 19%, 45%, and 26% of the cells, respectively. To a lesser extent, Bcat1 was also expressed by fibroblasts (CRC, OvC, and BC), dendritic cells (CRC, BC, and LC), and tumor cells (OvC). In general, Bcat2 showed expression trends similar to the ones seen by Bcat1 with a few exceptions such as a typically lower expression of the Bcat2 gene in myeloid cells and a preferential expression of Bcat2 gene by tumor cells of colon, breast, and lung origin (Figs. 6A, 6B and S4A).

**Figure 6.**
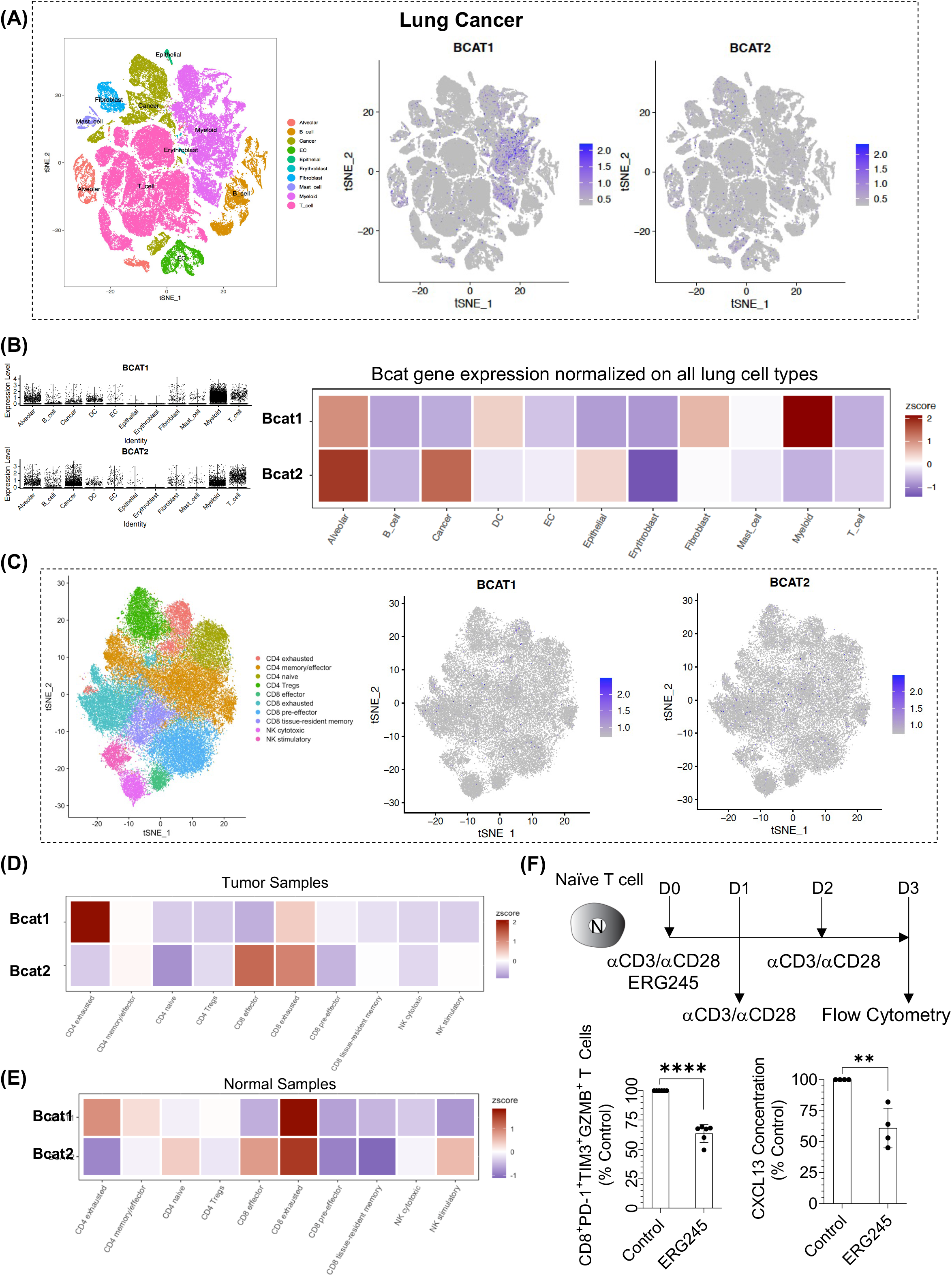
(A) Bcat1 and Bcat2 expression in various cell type clusters as visualized from sc-RNAseq data obtained from 8 lung cancer patients (50). (B) Summary of Bcat1 and Bcat2 expression in the cell clusters of (A). (C) Bcat1 and Bcat2 expression in T cell sub-populations, found in >36,000 T cells pooled from 8 lung cancer patients. (D) Heat map comparing gene expression of BCAT1 and BCAT2 proteins in the various T cell sub-populations found in the TME of lung cancer patients. (E) Heat map comparing gene expression of BCAT1 and BCAT2 proteins in the various T cell sub-populations from normal tissue samples excised near tumors of lung cancer patients. (F) Effect of ERG245 treatment on the phenotype of terminally exhausted CD8^+^ T cells, generated in vitro through repeat cycles of αCD3/αCD28 activation in the presence of TGF-β. From top down and left to right: Depiction of the experimental design, frequency of CD8^+^PD-1^+^TIM-3^+^GZMB^+^ T cells in the absence and presence of ERG245, and extracellular levels of cytokine CXCL13, secreted after 3 cycles of activation; 4 individual donors.

Lastly, we determined the heterogeneity of Bcat1 expression within the various T cell subgroups as clustered by Qian et al (41). We focused on T cells from tissues of LC patients because T cells constituted the most populous cell cluster in those samples (Fig. 6A). T cell populations exhibited relatively low frequency and low expression levels of Bcat1 and Bcat2 genes (Fig. 6C). Qualitatively, Bcat genes appeared to be expressed preferentially by intratumoral exhausted CD4^+^ and CD8^+^ T cells (Figs. 6D, S4B, and S4C) with some preference of intratumoral exhausted CD4^+^ T cells for Bcat1 and intratumoral exhausted CD8^+^ T cells for Bcat2. Statistical analysis using the Wilcoxon signed-rank test confirmed that BCAT1 was significantly more expressed in CD4^+^ and CD8^+^ exhausted T cells compared to the other T cells subtypes (p<0.0001 and p<0.001, respectively). Association of Bcat expression with T cell exhaustion was also seen in T cell populations of normal lung tissues, obtained from areas adjacent to the tumors, although, in that case, exhausted CD8^+^ T cells preferentially expressed Bcat1 (Fig. 6E). To examine if expression of Bcat1 is important in the differentiation of CD8^+^ T cells to terminally exhausted T cells (Tex), we generated Tex in vitro by subjecting naïve hCD8^+^ T cells, isolated from the peripheral blood of 4 donors, to successive cycles of αCD3/αCD28 activation in the presence of TGF-β and determined their phenotype after exposure to ERG245 (Fig. 6F). In the scRNA-seq dataset of Ref. 41, terminally exhausted CD8^+^ T cells were identified based on the presence of cytotoxic genes (Gzma, Gzmb, Gnly, Ifng), inhibitory genes (Pdcd1, Lag3, Tigit), and Cxcl13 expression (Fig. S3D). Repeat activations in the absence of TGF-β, which is the typical methodology for generating Tex in vitro (43), gave rise to CD8^+^ T cells that co-expressed cytotoxic and inhibitory molecules but did not secrete CXCL13. The latter required the addition of TGF-β to the cell media during cell activation (44). When hCD8^+^ T cells were exposed to ERG245 during the first cycle of activation but not in any of the subsequent cycles, then a decrease in the frequency of CD8^+^PD-1^+^Tim3^+^GZMB^+^ T cells was seen (Fig. 6F) along with a concomitant decrease in the extracellular levels of CXCL13 (Fig. 6F).

## DISCUSSION

To our knowledge, this is the first study to explore the function of BCAT1 in CD8^+^ T cell biology and to report on the unique role of the protein in the biology of the cells, a role that is mediated through a non-canonical mechanism of action involving restriction of mt iron levels and repression of OXPHOS. In contrast to what was reported for the role of BCAT1 in CD4^+^ T cell activation by Ananieva et al (45), BCAT1 does not appear to influence BCAA levels in CD8^+^ T cells nor does it participate in a feedback mechanism that restricts mTORC1 activation by metabolizing Leu. Yet, our data suggest that BCAT1 is required for the in vitro differentiation and in vivo priming of naïve CD8^+^ T cells to fully functional effector cells. Abrogation of IFNγ and GZMB expression was seen with BCAT1i in hCD8^+^ T cells, whereas mCD8^+^ T cells isolated from bcat1 KO animals showed suppressed levels of cytotoxicity following activation with αCD3/αCD28 antibodies compared to age-matched wt counterparts. Furthermore, systemic BCAT1i in animals, suffering from DSS-induced colitis, severely suppressed the number of activated CD8^+^ T cells found in MSN LNs and reversed the clinical manifestation of the disease suggesting that BCAT1i might be a novel therapeutic strategy for the treatment of autoimmune conditions, driven by hyperactive CD8^+^ T cells.

Even though functional BCAT1 appeared essential in the differentiation of the CD8^+^ T cells towards an effector phenotype, scRNAseq data indicated that effector cells, present in normal tissues as well as in tumor specimens of human cancer patients, possessed minimal Bcat1 gene expression compared to other T cell subpopulations; maximal Bcat1 gene expression was observed with exhausted CD4^+^ and CD8^+^ T cells. To reconcile the above observations, we considered the data collected from experiments interrogating the mechanism of action of BCAT1. The experiments suggested that two major events occurred following BCAT1i in activated CD8^+^ T cells: an early increase in mt iron levels accompanied by an increase in OXPHOS and an intracellular accumulation of lactate. Furthermore, OXPHOS appeared amplified long after BCAT1i was withdrawn indicating that BCAT1i induced irreversible metabolic reprogramming of CD8^+^ T cells, possibly through epigenetic changes linked to an altered iron homeostasis. On the other hand, intracellular levels of lactate were normalized soon after ERG245 was removed from the culture of the cells. Since Complexes I, II, and III of the electron transfer chain (ETC) require numerous Fe-S clusters for proper function, the increase in OXPHOS following the increase in mt iron is not surprising. Considering that an impaired OXPHOS was recently linked to CD8^+^ T cell exhaustion following chronic antigen stimulation (18), it is conceivable that BCAT1 participates in metabolically rewiring the T cells towards an exhausted phenotype by restricting mt [Fe-S] cluster biogenesis and limiting OXPHOS. In that light, BCAT1i is expected to prevent and/or suppress the rise of T_ex_ cells in the TME, while increasing the presence of T_eff_ cells. In support of the above hypothesis, we found an increased number of CD8^+^GZMB^+^ T cells in the TME of CT26 tumors in mice treated with a combination of ERG245 and an αPD1 antibody compared to corresponding tumors from mice receiving only αPD1 therapy (Fig. 2D). At the same time, BCAT1i suppressed CD8^+^ T cell exhaustion, generated in vitro through successive cycles of activation in the presence of TFG-β (Fig. 6F).

Although the link between BCAT1, mt iron, and OXPHOS provides a mechanistic explanation for the potential role of BCAT1 in T cell exhaustion, the mechanism of action of BCAT1 during differentiation of naïve CD8^+^ T cells towards effector cells remains unclear. The increased OXPHOS, observed following BCAT1i, is generally expected to increase the effector functions of the cells, especially since it does not affect aerobic glycolysis (46). Nonetheless, the opposite occurs. Bulk RNAseq analysis of control CD8^+^ T cells and CD8^+^ T cells treated with ERG245 for 24h revealed that BCAT1i is associated with a decrease in major cellular processes such as ribosome biogenesis, DNA replication, and mRNA splicing (Supplementary Table S3 and Figs. 4C and 4D), processes that are known to depend on [Fe-S] cluster proteins such as ABCE1 (which controls ribosome maturation and/or translation initiation) and DNA primase (which is required for DNA replication) (47,48). It is thus conceivable that the demand in mt [Fe-S] cluster biosynthesis, needed to support the increase in OXPHOS after BCAT1i, limits the availability of [Fe-S] clusters elsewhere in the cells. Alternatively, BCAT1i initiates a cascade of events that restricts iron-dependent mechanisms in the nucleus. Other mechanisms potentially contribute to the suppression of effector functions including accumulation of intracellular lactate and a possible rise in oxidative stress, usually associated with increased OXPHOS (49). Lactic acid is known to have deleterious effects on CD8^+^ T cell activation, whereas the dramatic increase in the humanin-like genes, seen in Fig. 4A, corroborates the presence of oxidative stress in the cells following BCAT1i (50). Taken together, the above observations suggest that BCAT1 plays a major regulatory role in iron homeostasis, which, in activating CD8^+^ T cells, ensures proper ribosome biogenesis, DNA replication, and adoption of a cytotoxic phenotype. Under conditions of chronic antigen exposure, BCAT1 contributes to the reprogramming of the cells towards exhaustion.

Our data additionally highlight the potential of BCAT1 as a druggable target in immuno-oncololgy and support the idea of pulsing BCATi as an immuno-tuning tool to increase cytotoxicity of tumor infiltrating lymphocytes (TILs) in the TME. In our hands, aspects of the phenotype of CD8^+^ T cells that emerged following BCAT1i (lack of effector functions, downregulation of genes associated with ribosome biogenesis) were completely reversed upon ERG245 withdrawal, while other aspects were sustained and even enhanced (OXPHOS). The combined effect was the rise of super energetic CD8^+^ T cells with increased cytotoxicity and higher potential for cell growth due to increased ribosome biogenesis. We have speculated that the appearance of those CD8^+^ T cells is the result of epigenetic changes driven by certain histone modifications such as downregulation of the histone acetyltransferase Kat2b (which is expected to increase ribosome biogenesis) and downregulation of histone methyltransferase Kmt2a (which is expected to enhance OXPHOS) (Fig. 5E) (51,52). As an immuno-tuning tool, BCAT1i can be used either in a CAR-T cell setting, prior to the cell transfer to prevent and/or suppress the rise of an exhaustive phenotype in the CAR-T cells, or in combination with an ICI, to increase the cytotoxicity of reinvigorated CD8^+^ T cells in the TME. Our in vivo experiments provide evidence for both strategies. Brief administration of ERG245 in combination with αPD1 therapy dramatically increased the efficacy of the αPD1 resulting in extensive number of complete tumor regressions. ACT transfer of ERG245-treated CD8^+^ T cells increased survival of CT26 tumor-bearing mice compared to untreated CD8^+^ T cells. Although not exceptional at first glance, the data of the Fig. 2E become promising if one considers that the ACT experiments involved the transfer of a relatively low number of CD8^+^ T cells per animal and that the transferred cells were not engineered to recognize cancer cells. As a result, a very small fraction of transferred CD8^+^ T cells likely reached the tumor site and even a smaller penetrated into the TME, where it likely became rapidly hypofunctional. This is the reason why the ACT of untreated CD8^+^ T cells had no effect on tumor growth despite the co-administration of αPD1. Collectively, our data suggest that BCAT1i provides a unique opportunity to formulate therapeutic strategies for BCAT1 inhibitors to be used for both dampening and enhancing CD8^+^ T cell cytotoxicity.

Finally, the study provided evidence that, even though BCAT1 enzymatic activity is essential for mt iron homeostasis and OXPHOS regulation, BCAT1 does not appear to play a role in controlling intracellular levels of BCAAs nor does it appear to contribute to Glu synthesis. Subsequently, it is reasonable to assume that the mechanism of action of BCAT1 in CD8^+^ T cells is not mediated via fluctuating levels of metabolites such as BCAAs or αKG but rather involves the protein itself. Here, we hypothesize that BCAT1 is part of a mechanism mediated via the IRP1/ACO1 protein and designed to meet the iron demands of the CD8^+^ T cells at the subcellular level after αCD3/αCD28 activation. A schematic of the proposed mechanism is presented in Fig. 7. Briefly, iron is an essential micronutrient, acting as a cofactor for [Fe-S]-cluster containing proteins and heme (53). Activated T cells exhibit increased need for iron utilization and, in response to TCR engagement, T cells rapidly translocate their transferrin receptor TfR1 from the endosomal compartment to the immunological synapse to ensure proper iron levels. At the same time, defective iron transport is linked to a global defect in T cell proliferation and combined immunodeficiency (54-56). Because iron dysregulation can lead to oxidative damage, intracellular levels of iron are tightly controlled through the actions of two iron regulating proteins: cytosolic IRP1 and mitochondrial IRP2 (39). When the iron levels in the cells are low, IRPs initiate the translation of proteins involved in iron transport and storage. Under replete conditions, IRP1 acquires a [4Fe-4S] cluster, which functions as a molecular switch and converts the protein to ACO1, the enzyme responsible for the stereo-specific conversion of citrate to isocitrate via cis-aconitate in the cytoplasm (57). The actions of the two states of IRP1 (also known as apo- and holo-IRP1) are considered mutually exclusive. In an intriguing study, Huynh et al reported that the *Drosophila* analog of IRP1, in its holo-state, translocates to the nucleus, where it dowregulates iron-dependent processes (58). The translocation was cell-type specific and required binding with the *Drosophila* analogs of glycogen branching enzyme (GBE1) and CISD1. The investigators indicated that, in the nucleus, holo-IRP1 interacted with ribosomal proteins, eukaryotic factors, and histone proteins and proposed that holo-IRP1 acts as a sensor to restrict iron-dependent processes in the nucleus once peak demand for iron has passed. They additionally provided evidence that human ACO1 exhibits similar behavior under specific circumstances, an observation that was corroborated by Gu et al in human hepatoma cell lines (59). These reports complement our data, which suggest that BCAT1 prevented the iron-sulfur switch from the apo- to the holo-state of IRP1 by forming a complex with the former. BCAT1i increased ACO1 activity in Jurkat and hCD8^+^ T cells, indicating a successful transition to the holo-state. In CD8^+^ T cells, BCAT1i was accompanied by transcriptomic dowregulation of ribosome biogenesis, cholesterol biosynthesis, mRNA translation, and DNA metabolism, all iron-dependent nuclear processes. We speculate that BCAT1i-driven downregulation of the above nuclear pathways were the results of a successful translocation of ACO1 to the nucleus. Subsequently, we hypothesize that BCAT1 is essential for CD8^+^ T cell differentiation because it allows the cells to apply the ACO1 “break” in the nucleus in a timely fashion. Conversely, BCAT1i forces the cells to apply the ACO1 “break” too early leading to abrogation of effector functions. Withdrawal of BCATi releases the ACO1 “break” and allows the nuclear processes to proceed undeterred in cells that, in the meantime, have acquired a superior energetic status and, thus, the potential for increased cytotoxicity.

**Figure 7.**
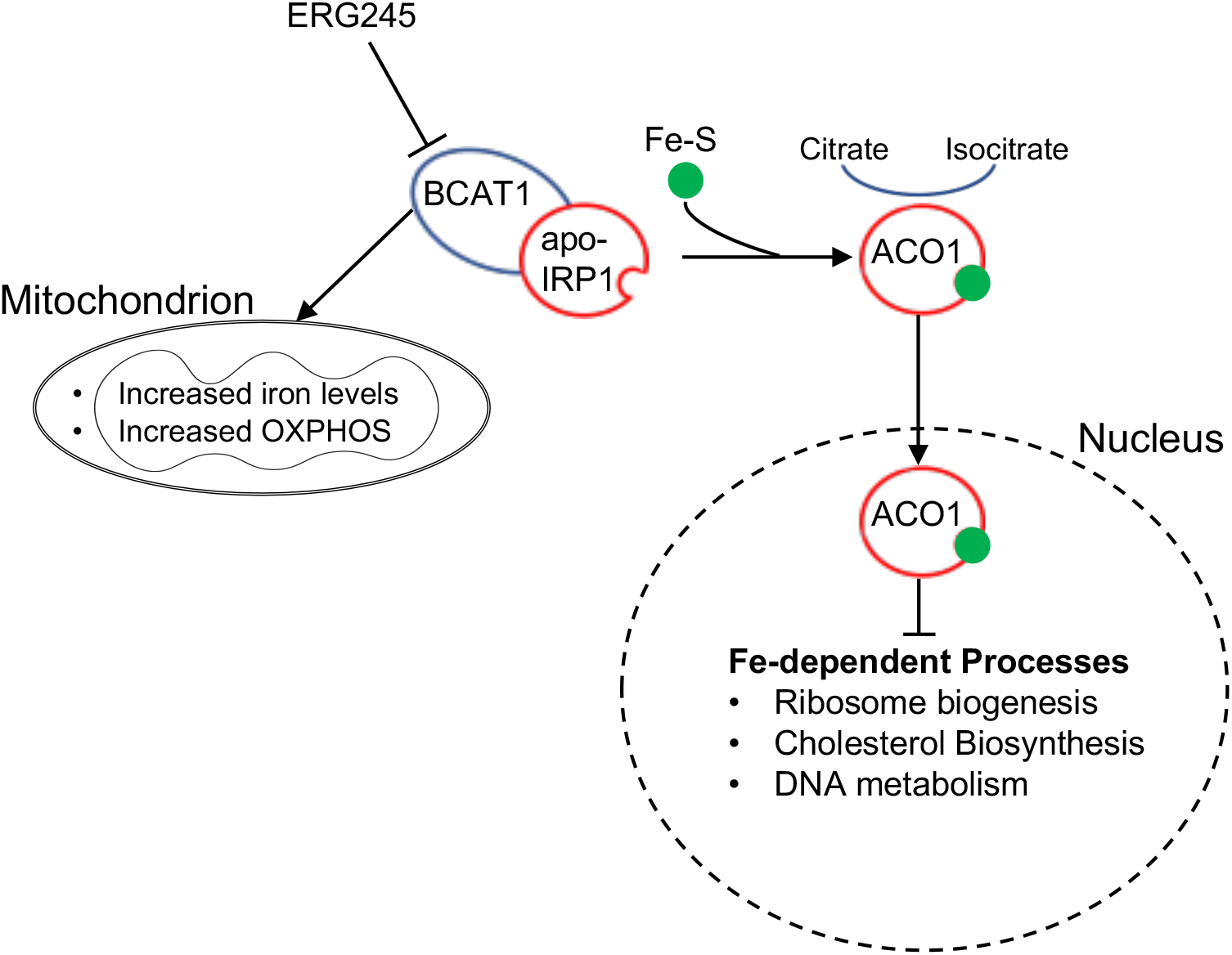
Schematic of the proposed mechanism of action of BCAT1i. ERG245 inhibits BCAT1 and allows apo-IRP1 to pick up Fe-S clusters and convert to ACO1, which then translocates to the nucleus of CD8^+^ T cells and restricts iron-dependent processes leading to abrogation of effector functions.

## MATERIALS AND METHODS

### ERG245 Dosing

For FACS experiments analyzing expression of cytotoxic markers at 24h hCD8^+^ T cells were treated with 200 μM ERG245, which greatly reduced the frequency of IFNγ- and GZMB-expressing. The same concentration was utilized for RNAseq experiments with cells treated with ERG245 for 24h. Because treatment of CD8^+^ T cells with 200 μM ERG245 leads to cell apoptosis (upon withdrawal or persistent exposure beyond 24h), a lower concentration (100 μM ERG245) was utilized for experiments lasting more than 24h, such as for proliferation assays or for in vitro experiments requiring temporal BCAT1i. All Seahorse experiments were performed with 100 μM ERG245 in order to minimize cell death. For metabolic experiments, CD8^+^ T cells were treated with 200 μM ERG245 and wt Jurkat cells were treated with 400 μM ERG245, as this concentration was shown to be the IC_50_ value for inhibition of Jurkat cell proliferation. For IP experiments involving rhBCAT1, 10 nM ERG245 were used. This concentration mirrors the inhibitory concentrations used in the enzymatic assays with rhBCAT1. DSS-colitis testing and in vivo tumor growth experiments involved dosing of the animals with 5 mg/kg ERG245 twice a day. According to PK analysis, a 5 mg/kg ERG245 dose is expected to produce a max circulating concentration in the 100-200 mg/kg range.

### Cell Isolation

hCD8^+^ T cells were isolated from commercially available sources (buffy coats from ZenBio Inc. and leukopaks from Miltenyi Biotec) using magnetic separation (positive selection; all microbead kits from Miltenyi Biotec) and the MultiMACS Cell24 Separator. Single cell suspensions from murine MSN LNs and spleens were obtained using a gentleMACS Dissociator (Miltenyi Biotec) and published protocols by Miltenyi Biotec. Purified mCD8^+^ T cells from spleens were obtained using magnetic separation (negative selection). For ACT experiments, spleen-derived mCD8^+^ T cells were further purified using a FACSAria cell sorter (BD Biosciences). Single cell suspensions from CT26 tumors were obtained using a tumor dissociation kit from Miltenyi Biotec per manufacturer’s instructions.

### CD8^+^ T Cell Culture

Purified hCD8^+^ or mCD8^+^ were activated using Dynabeads from ThermoFisher Scientific per manufacturer’s instructions or using plates coated with αCD3 antibody. In the latter case, the appropriate number of wells of a flat-bottomed tissue culture plate were coated with 10 μg/mL αCD3 (Biolegend) in PBS for 3h at 37°C. The wells were then washed with PBS 2X. After addition of CD8^+^ T cells, 5 ug/mL αCD28 antibody were also added into the cultures. hCD8^+^ T cells were cultured in RPMI (ATCC modification) containing 10% FBS and antibiotics. rhIL-2 (30U/mL, PeproTech) was typically included in the culture of hCD8° T cells 24 hrs after activation. In contrast, mCD8^+^ T cells were cultured in RPMI (ATCC modification) containing 10% FBS, antibiotics, and 2 mM glutamine, 50 μM mercaptoethanol, 0.5 ng/mL IL-7, 30U/mL IL-2, and 1% Insulin Transferrin Selenium-A (ITS) (complete murine RPMI media). To exhaust CD8^+^ T cells, purified hCD8^+^ T cells were subjected to 3 consequent cycles of αCD/αCD28 activation in the presence of 50 ng/mL TGFβ (R&D Systems) and 30U/mL rhIL-2 lasting 24h each.

### Jurkat Cell Culture

Jurkat BCAT1^-/-^ cells were generated by electroporation of pre-assembled sgRNA-Cas9 RNPs according to Ref (Lin 2014) using in vitro transcribed (IVT) sgRNAs and purified Cas9-NLS. Details are provided in Supplementary Methods. Jurkat BCAT1^-/-^ cells (Bcat1 KO), Jurkat cells generated using a non-targeting RNP carrying a control sgRNA (Control KO) and parental Jurkat (wt) were cultured in RPMI containing 10% FBS and antibiotics.

### BCAT1 Enzymatic Assays

The assays performed essentially as described in Ref. 61. The assays used 15 μg LeuDH from EMD Millipore and 200 ng BCAT1 or BCAT2 from Abcam. Enzymatic reactions rates were measured every min over a period of 10 min at 37°C using a Molecular Devices SpectraMax M2 microplate reader.

### Aconitase Activity

To determine the aconitase activity of in the cytosol of transfected Jurkat cells, 3×10^6^ cells were activated for 3h and lysates were obtained as described in the Technical Bulletin accompanying the Aconitase Assay Kit from Sigma. Aconitase activity in the lysates is determined per manufacturer’s instructions. To determine the aconitase activity of rhIRP1 (Origene or Abcam), 1 μg hrBCAT1 (Abcam or Abnova) was mixed with 0.5 μg hrIRP1 (Origene or Abcam) in 50 uL of reaction buffer (Aconitase Assay Kit from Sigma). After addition of 10 μL activating solution ((NH_4_)Fe(SO_4_)_2_ and cysteine; Aconitase Assay Kit from Sigma), the reaction mixture was incubated at 37°C for 1h and its aconitase activity was determined per manufacturer’s instructions (Aconitase Assay Kit from Sigma).

### Flow Cytometry Analysis

For analysis of intracellular biomarkers of purified cells, CD8+ T cells were activated overnight as previously described, brefeldin (Biolegend) and monensin (Biolegend) were then added to the cultures, and the cells were additionally incubated for 4 hrs at 37°C and 5% CO_2_. The cells were subsequently collected, washed with PBS and stained with Zombie Violet fluorescent dye (Biolegend) per manufacturer’s instructions for live/dead discrimination. After removal of the dye, the cells were re-suspended in flow cytometry staining buffer (PBS containing 4% FBS and 2 mM EDTA) and blocked with TruStain FcX (mouse or human, Biolegend) for 5 min at 4°C. Subsequently, the cells were exposed to a cocktail of antibodies against extracellular biomarkers (if any) for 30 min at 4°C, washed with staining buffer 2X, and fixed using 2% formaldehyde (10 min at 4°C). At the end of the fixation period, the washing step was repeated. If the experiment included intracellular biomarkers, the cells were permeabilized using a permebilization buffer (Invitrogen), stained with a cocktail of antibodies against the intracellular biomarkers (30 min at 4°C), washed with the permeabilization buffer 2X and resuspended in staining buffer. Biomarker analysis was performed with BD FACSCanto II. The experiment typically included appropriate isotype control sample(s). Data were analyzed using FlowJo v10.

### CD8^+^ T cell proliferation

Cell proliferation of hCD8^+^ T cells were assessed using carboxyfluorescein succinimidyl ester (CFSE), a membrane-permeable green fluorescent label. Briefly, naïve CD8^+^ T cells were incubated with 5 μM CFSE in PBS for 20 min at 37°C. The CFSE solution was then removed, and the cells were washed with PBS containing 10% FBS. The cells were subsequently activated and cultured as previously described for 72-96 hrs. The cells were then collected and analyzed by flow cytometry, as previously described. The number of cells per cell division was estimated using FlowJo. The Weighted Proliferation Index was estimated using the following formula: (0*P_0_+1*P_1_+…+N*P_N_)/((N*(N+1)/2), where P_x_ is the percentage of cells that have undergone x number of divisions and x is a number between 0 (undivided cells) and N (maximal number of cell divisions observed).

### Metabolomic Studies

In a typical experiment, 1-2×10^6^ hCD8^+^, mCD8^+^ or Jurkat cells were activated with αCD3/αCD28 antibody cocktail in complete RPMI SILAC media comprising of RPMI SILAC, 48 mg/L L-Lysine hydrochloride, 200 mg/L L-arginine, 2 g/L sodium bicarbonate, and 50 mg/L ^15^N-Leucine in the presence or absence of 200 μM ERG245 for 24h (h/mCD8^+^ cells) or 400 μM ERG245 for 3h (Jurkat cells). At the end of the incubation period, conditioned media and cells were collected. The cells were washed 2X with Tris buffer pH 8.0 and lysed/fixed in 400 μL methanol, previously chilled at −20°C. After addition of 400 μL ice-cold ultrapure water and 4 μg 2,2,3,3 D_6_-succinic acid as internal standard (IS), 400 μL dichloromethane, previously chilled at −20°C, were added to the samples. The samples were vortexed briefly and centrifuged at 12,000 g (10 min, 4°C) using a TOMY High Speed Refrigerated Micro Centrifuge (TX-160). The water-soluble layer of each sample was transferred into a clean glass vial and then dried under vacuum at 4°C using a refrigerated CentriVap Concentrator (Labconco). Conditioned media were processed similarly with the exception that 4 μg of internal standard were added into 200 μL of conditioned media and then the media were mixed with 200 μL methanol and 200 uL dichloromethane. Water soluble organics, present in the dried samples, were derivatized using N-methyl-N-trimethylsilyl trifluoroacetamide (MSTFA) and analyzed using GC-MS. Details are provided in the Supplementary Methods.

### Bioenergetics

Oxygen consumption rate, extracellular acidification rate, and ATP rate were determined by using a Seahorse XF96 Extracellular Flux Analyzer (Agilent). Briefly, one hour before the experiment, hCD8^+^ T cells were resuspended in unbuffered, serum-free DMEM media and plated on 96-well Seahorse plates coated with Cell-Tak (BD Biosciences, San Jose, CA) at 250,000 cells/well. Cells were centrifuged at 200 g for 5 min and incubated into a 37°C non-CO_2_ incubator for 45 minutes prior to assay. Measurements were performed after injection of compounds affecting bioenergetics: 1 μM oligomycin, 0.5 μM carbonyl cyanide 4- (trifluoromethoxy)phenylhydrazone (FCCP), 1 μM Rotenone, 10 mM glucose and 50 mM 2-Deoxy-D-glucose (2-DG). Upon completion of the Seahorse XF96 Flux measurements, the results were collected through the Seahorse XF Stress Test Report Generator, and analyzed.

### Metal Analysis

In a typical experiment, 10-20×10^6^ Jurkat cells (wt, Control KO, Bcat1 KO) were activated in 60 mm^2^ tissue culture dishes (plated with αCD3 antibody) for 1h; αCD28 antibody was also add to the cultures. At the end of the incubation period, cells were collected and washed in PBS 1X, then in ice-cold salt-free iso-osmotic buffer A (300 mM sucrose, 10 mM Tris and 1 mM EDTA adjusted to pH 7.4 with 85% phosphoric acid), and finally in ice-cold salt-free iso-osmotic buffer A (300 mM sucrose, 10 mM Tris and 1 mM EDTA adjusted to pH 7.4 with glacial acetic acid). Mitochondria from washed Jurkat cells were obtained using a Mitochondria Isolation kit (Boster Biological Technology), whereas nuclear fractions were obtained using a Cytoplasmic and Nuclear Protein Extraction kit, (Boster Biological Technology) per manufacturer’s instructions. Purified mitochondria or nuclear proteins were acidified with 70% HNO_3_, OPTIMA Grade and stored at −20C until ICP analysis. Prior to ICP analysis, samples were diluted 1:70 with ultra-pure water to a final concentration of 1% HNO3, divided into 3 fractions, spiked with 2,4, and 6 ppb Fe, respectively, and analyzed using an iCAP 7000 Series ICP-OES (ThermoFisher Scientific). A calibration curve was also generated. The reportable linear range of the assay was also determined along with the lower limit of quantification. Each experiment was performed in triplicate and repeated 5 times.

### Immunoprecipitation Assays

1 μg of rhBCAT1 tagged with GST (Abnova) was transferred °into 100 μL reaction mixture solution, recommended for use in BCAT1 enzymatic assays (61), in the presence or absence of 10 nM ERG245. The reaction mixture was allowed to stand at 37°C for 15 min. Subsequently, 0.5 μg of rhACO1 tagged with His (Abcam) was added to the reaction mixtures and the samples were rotated for 30 min at 37°C. At the end of the incubation period, 1 μg and anti-GST (GeneTex) or anti-His (Abcam) were added to the samples. One sample received an irrelevant antibody (a mouse IgG1k against IFNγ from Biolegend) control. The samples were rotated at room temperature for 1 h. Protein-antibody complexes were immunoprecipitated by rotating the samples with 40 μL protein A-agarose beads (ThermoFisher Scientific) overnight at 4°C. The samples were then washed with 2X with TBS containing 0.5% Tween-20 and immunoprecipitated proteins were obtained by heating the samples in 25 μL Laemmli buffer containing 5% mercaptoethanol at 95°C for 5 min. Immunoprecipitation of rhBCAT1 and rhIRP1 proteins was also performed in the presence of a (NH_4_)Fe(SO_4_)_2_/cysteine solution with and without ERG245. Briefly, 1 μg rhBCAT1 (Abcam) was incubated with ERG245 in 50 μL of reaction buffer from the Aconitase Assay Kit (Sigma) for 15min at 37°C. Subsequently, 0.5 μg rhIRP1 (Origene) was added to the reaction mixture along with 10 μL of (NH_4_)Fe(SO_4_)_2_/cysteine solution (Aconitase Assay Kit, Sigma). The mixture was incubated for 30 min at 37°C. In this case, hrBCAT1, which is His-tagged, was immunoprecipitated with an anti-His antibody (Abcam) and the presence of rhIRP1 was detected with an anti-IRP1 antibody (Abcam).

### Western Immunoblotting

Approximately, 1×10^5^ hCD8^+^ T cells were heated in Laemmli sample buffer for 2 min at 90°C and then sonicated (3 pulses, 5s each). Cell lysates were heated for an additional 4 min at 90°C and sonicated again. Protein present in T cell lysates were resolved by SDS-PAGE, transferred onto nitrocellulose membranes, and immunoblotted using appropriate primary and secondary antibodies. The probed proteins were detected using SuperSignal West Femto chemiluminescent substrate (ThermoFisher Scientific).

### Quantitative PCR Analysis

Total RNA was extracted from 0.5×10^6^ naïve and activated CD8^+^ T cells using a using an RNeasy Mini kit (Qiagen) and additionally purified using an RNase Free DNase Set (Qiagen) according to the manufacturer’s instructions. Quantitative conversion of RNA to single-stranded DNA was executed using a high Capacity cDNA Reverse Transcription kit (Applied Biosystems). Reaction mixtures were prepared using FastStart TaqMan Probe Master Mix (Applied Biosystems). All quantitative PCR reactions were performed using a QuantoStudio 6 Flex (Applied Biosystems) and the following conditions: Hold (20s@95°C) and Cycle (40 cycles: 1s@95°C, 20s@60°C). Results were analyzed using the comparative Ct method, and each sample was normalized to the reference mRNA level of the *B2M* gene.

### RNAseq Analysis

Total RNA was isolated and DNase I treated using the RNeasy Plus Mini Kit from Qiagen. RNA integrity was checked by Agilent Bioanalyzer 2100 or Agilent Tapestation 4200 and samples with clean rRNA peaks (RIN>7) were used for library preparation. RNA-Seq library was prepared according to KAPA mRNA HyperPrep kit with 200-300 bp insert size (Roche) using 250 ng of total RNAs as input. Final library quality and quantity was analyzed by Agilent Bioanalyzer 2100 or Agilent Tapestation 4200 and Qubit 3.0 Fluorometer from Thermo Fisher Scientific Inc. 150 bp PE (paired end) reads were sequenced on Illumina HiSeq X sequencer. Bioinformatics studies are described in the *Supplementary Text*. RNA

### In Vivo Experiments

#### PK Analysis

The study was executed by Pharmacology Discovery Services Taiwan Ltd, a subsidiary of Eurofins (Item No: 515570) US. The work was performed in accordance with the Guide for the Care and Use of laboratory animals (National Academy Press, Washington, DC, 2011) using an AAALAC accredited facility. The study protocol was approved by the Pharmacology Discovery Services IACUC. Briefly, non-fasting pharmacokinetic parameters were obtained after administration of 1 mg/kg i.v. or 10 mg/kg p.o ERG245 to C57BL/6 mice and blood collection at 3, 10, 30, 60, 120 and 240 min (i.v. dosing) or 10, 30, 60, 120, 240 and 360 min (p.o. dosing). Each animal was subjected to a single blood draw and 3 animals per time point were used. Plasma was also obtained from 3 naïve (drug-free) mice as blank controls. A calibration curve was generated by spiking drug-free samples with known concentrations of ERG245. Spiked samples and unknown samples were processed together using acetonitrile precipitation and then analyzed by LC-MS/MS. The reportable linear range of the assay was also determined along with the lower limit of quantification. ERG245 plasma concentrations were plotted against time. The fundamental PK parameters of ERG245 after iv or po dosing were determined from the non-compartmental analysis of the data using WinNonlin. The oral bioavailability was also calculated. Experimental details are provided in *Supplementary Text*.

#### Isolation of Spleens from Bcat1 KO Animals

*Bcat1*^-/-^ and wt littermate mice were previously described (31) and maintained according to protocols approved by the University of Virginia Animal Care and Use Committee. Spleens collected immediately following euthanasia and stored in tissue storage solution (Miltenyi). CD8^+^ T cell were isolated 24h later as previously described.

### Animals Models

Disease Animal models were conducted at Washington Biotechnology, Inc. The studies were approved by the local Institutional Animal Care and Use Committee (IACUC). The animal research facility is registered with the US Department of Agriculture, has an Animal Assurance Statement with the Office of Laboratory Animal Welfare (OLAW), and is AAALAC accredited.

#### Acute DSS-Induced Colitis Model

Female C57BL/6 mice (Charles River Laboratories, 18-20 g each) were received, individually examined and housed in cages of five mice each. The mice were placed in quarantine with daily inspections. The mice were ear-tagged with unique numbers for identification purposes, weighed, and sorted into treatment groups based on average body weight. The mice were then allowed *ad libitum* access to 3% wt/v DSS/water for the duration of the experiment. Treatment was initiated 24h after exposure of the mice to DSS. Mice were treated with either 5 mg/kg ERG245, ip, twice a day or with Vehicle control for a total of 6 days (n=4 mice/group). Mice were observed daily for clinical signs of disease and scored according to the following system: Stool consistency (<u>0</u>=separate hard lumps, <u>1</u>=sausage-like with cracks, <u>2</u>=soft blobs, <u>3</u>=watery, no solid pieces), Anal prolapse (<u>0</u>=not visible outside the body, <u>1</u>=visible outside the body), Fecal blood (<u>0</u>=no blood in the stool, <u>1</u>=blood in the stool), Weight loss (<u>0</u>= body weight loss≤1%, <u>1</u>= 1<body weight loss≤5%, <u>2</u>=5<body weight loss≤10%, <u>3</u>=10<body weight loss≤15%, 4= body weight loss>15%). Disease Index (DI) is defined as a sum of all scores. The study was terminated seven days after initial exposure to DSS using CO^2^-induced euthanasia. MSN LNs were removed and stored in tissue storage solution (Miltenyi Biotec) until they were processed for flow cytometry, typically within 2h from collection.

#### Syngeneic CT26 Tumor Model Studies

##### Survival Study

Female Balb/c mice 6-7 weeks were injected with 1×10^5^ CT26 tumor cells in PBS containing 20% matrigel in the right flank. Tumors were allowed to grow over a period of 7-14 days to a predetermined initial tumor volume. Mice were then randomized to the appropriate number of groups receiving a specific treatment regimen. The dosing regimen was initiated one day after group randomization. All mice received αPD-1 or isotype control antibody, both from BioX Cell, at 10 mg/kg on indicated days. ERG245 was administered at 5 mg/kg twice a day on indicated days. Mice were observed daily for clinical signs of stress. Tumor volumes and body weights were assessed every other day. Mice were terminated when tumor volumes reached 2000 mm^3^, as measured using a digital caliper and the following formula: ½(Length x Width^2^). Additional information regarding animal maintenance is provided in Supplementary Methods.

##### Adoptive T Cell Transfer Study

Female Balb/c mice were injected with CT26 tumor cells as previously described. When the tumors reached an average volume of ∼100 mm^3^, six female naïve Balb\c mice were then sacrificed and spleens were obtained and stored in tissue storage solution (Miltenyi) until use, typically within two hours of collection. Purified mCD8^+^ T cells were obtained as described in *Cell Isolation*. Subsequently, half of the cells were pre-exposed to 100 μM ERG245 in complete murine RPMI media (see above) for 1h at 37°C and 5%CO_2_. All cells were activated for 24h using Dynabeads (ThermoFisher Scientific) per manufacturer’s instructions. The following day, mCD8^+^ T cells from each group were counted and their viability was determined to be more than 97%. Tumor-bearing mice were randomized into two groups (ACT(untreated)) and (ACT(ERG245)) and then each mouse received via a tail-vein injection 2.75×10^6^ mCD8^+^ T cells. All the mice received 10 mg/kg αPD-1 at days 2, 5, 8, 11, and 14 after ACT.

### Statistical Analysis and Illustrations

Data were visualized and statistically analyzed using Student’s *t*-test (Microscoft Excel 2016 or Graphpad Prism 9) with the exception of Kaplan-Meier survival curves, where the logrank test was used (Graphpad Prism 9). P-values were typically reported as follows: *: *p<0*.*05*; **: *p<0*.*01*; ***: *p<0*.*001*. The Wilcoxon signed-rank test was used to compare Bcat1 gene expression in the various T cell sub-populations.

## Contributions

F.L. and D.L. performed the bioinformatics analysis of the scRNAseq data. M.C., H.E., and C.M. performed the Seahorse analysis. W.L. performed the synthesis of BCAT1 inhibitors. H.A.V. assisted with the flow cytometry experiments and the execution of the enzymatic assays. K.G. performed the WGCNA analysis. L.S. performed the iron analysis. I.L.P. assisted with the iron analysis and the study design. T.W., H.B., M.W,. C.Y., and J.E.C. generated the Jurkat Mock KO and Jurkat Bcat1 KO cells. K-S.P. with assistance from J-H.K. and J.B. provided the transgenic mice and necessary animal specimens. Additionally, JB provided constructive review of the manuscript. A.E.P. designed the study, supervised the execution of the experiments, organized the collaborations, review and analyzed the data, and wrote the manuscript.

