## Supplemental Material for "BCAT1 inhibition affects CD8^+^ T cell activation, exhaustion, and tumoral immunity by altering iron homeostasis"

#### Supplementary Text

**The Metabolomic Effect of BCAT1i on Lac and Glc.** Our metabolomic studies indicate that BCAT1i (hCD8$^+$ T cells) or Bcat1 gene deletion (mCD8$^+$ T cells) commonly increased intracellular levels of Lactate (Lac) as shown in Fig. 3B. At the same time, intracellular Glucose (Glc) levels increased in ERG245-treated hCD8$^+$ T cells compared to naïve and control cells. Increased uptake of Glc is a required event in the metabolic reprogramming of immune cells. While intracellular levels of Glc remained constant in mCD8$^+$ Bcat1 KO T cells compared to control cells, extracellular levels of Glc decreased with Bcat1 deletion indicating increased utilization of Glc (Fig. S1F). Interestingly, in both hCD8$^+$ and mCD8$^+$T cells, loss of BCAT1 function resulted in a slight decrease in extracellular levels of Lac indicating a possible impairment in the ability of the cells to exocytose Lac (Fig. S1F). Lac is an important metabolite, implicated in both suppression and augmentation of effector functions of TILs (S1,S2). Accumulation of endogenous Lac, present in the acidic form, is linked to a decrease in intracellular pH and complete suppression of CD8$^+$ cell cytotoxicity (S1). In contrast, exogenous Lac, present in a buffered form as in the case of Lac produced during exercise, can serve as a primary fuel during early differentiation of CD8$^+$ T cells resulting in effector cells with enhanced expression of GZMB and INF$\gamma$ (S1). It is thus conceivable that a build-up in intracellular Lac in lactic form contributes to the suppression of effector functions seen in ERG245-treated hCD8$^+$ and mCD8$^+$ Bcat1 KO T cells, whereas an extracellular build-up in a buffered lactate form might contribute to the increased effector functions seen upon withdrawal of BCAT1i. Interestingly, the effects of lactic acid on CD8$^+$ T cells are considered reversible once the source of lactic acid accumulation is eliminated. To determine if Lac build-up persists following withdrawal of ERG245, we activated CD8$^+$ T cells in the presence and absence of ERG245 for 24h in media containing U-$^{13}$C-Glc and collected conditioned media and cell lysates. An identical experiment was also performed, in which cells were washed at 24h and then incubated in media containing U-$^{12}$C-Glc and IL-2 and devoid of ERG245. In that experiment, conditioned media and cell lysates were collected at 48h. Isotopic analysis of Lac present in conditioned media (CM) and cell lysates (CL) revealed that (m+3)/(m+0) Lac ratios were higher in ERG245-treated cells compared to control cells at 24h confirming that, as activated cells replace intracellular U-$^{12}$C-Glc with U-$^{13}$C-Glc, (m+3) Lac accumulates intracellularly in the ERG245-treated cells thus raising the (m+3)/(m+0) ratio (Fig. S1G). As (m+3) Lac builds up intracellularly in the ERG245-treated cells, less (m+3) Lac is released and the isotopic (m+3)/(m+0) ratio of extracellular Lac is decreased in the ERG245-treated cells compared to control cells (Fig. S1G). When media were replenished with media containing U-$^{12}$C-Glc, intracellular (m+3)/(m+0), Lac ratios were diminished indicating that accumulated (m+3) Lac was readily exocytosed and/or metabolized (Fig. S1D). At 48h, control and ERG245-treated cells had similar (m+3)/(m+0) Lac ratios in both CM and CL suggesting that withdrawal of BCAT1i normalized Lac metabolism (Fig. S1H). The data collectively suggest that suppression of effector functions with BCAT1i is not the result of the failure of the cells to glycolytically reprogram their metabolism. Although a potential decrease in intracellular pH due to the presence of lactic acid might contribute to the repression of the effector phenotype, RNAseq data suggest a different mechanistic pathway.

**Materials and Methods**.

**PK Analysis**. C57BL/6 mice were subjected to bolus injections of 1 mg/kg ERG245 i.v. or 10 mg/kg ERG245 p.o. At predetermined time points, mice were sedated under general inhalant anesthesia (3 % isoflurane) and 300-400 µL of blood were collected in tubes coated with lithium heparin by cardiac puncture. The plasma was harvested after centrifugation and kept frozen at - 80°C until further processing. Proteins present in the plasma were precipitated by acetonitrile. Specifically, 50 µL of plasma samples were transferred to a 96-well plate and 150 µL of 0.1 ng/µL of oxybutyrin in acetonitrile were also added as internal standard. After sample mixing and centrifugation, 150 µL of supernatant were withdrawn from each well and mixed with 150 µL of water/formic acid 100/0.2 v/v for LC-MS/MS analysis. The samples were analyzed using an AB SCIEX API 3000 (LCMS-010) (ionization mode: electrospray, positive ions) and a Eurospher II 100-5 C18 column (gradient elution with two mobile phases: acetonitrile/formic acid 100/0.2 v/v and water/formic acid 100/0.2 v/v). The column temperature was 30°C, the injection volume was 10 µL, and the run time of the method was 5 min. The linear range of the assay was 10-10000 ng/mL. The lower limit of quantification was 10 ng/mL.

**Metabolite Quantification Using GC-MS Analysis**. A pyridine solution of methoxamine hydrochloride (MOX) was prepared at 40 mg/mL and allowed to stand for at least 30 min at room temperature. Subsequently, 20 µL of MOX solution were added into the dried samples. The samples were allowed to stand for 90 min at room temperature. After addition of 80 µL of MSTFA, samples were heated at 60°C for 60 min. At the end of the heating period, 80 µL of derivatized materials were transferred into a clean vial containing a 100 µL polypropylene insert and the samples are analyzed using a 7890A GC System (Agilent Technologies) equipped with an Agilent J&W DB-5ms column (30 m, 0.25 mm, 0.25 µm), connected to an 5975C Triple-Axis MS Detector (Agilent Technologies) and outfitted with an autosampler. The MS source was held at 230 °C and the quadrupole at 150 °C. The GC oven temperature was held at 50 °C for 5 min, increased to 240 °C at a rate of 10 °C $min^{-1}$, held at 240 °C for 5 min and increased to 300 °C at a rate of 10 °C $min^{-1}$. A volume of 1 µl was injected in the system in a 1:5 split mode. The amount of organics present in the sample was determined after peak integration at a specific m/z ratio. Metabolite levels were normalized using the IS.

**Generation of Bcat1 KO Jurkat Cells Using Electroporation**. Jurkat cells (from ATCC) were transfected with two RNPs in order to generate a deletion of ~214 bp across the border between exon 3 and the following intron (3-i), covering all relevant isoforms. The sequences of sgRNA templates for IVT and corresponding primers for PCR validation and Sanger sequencing are given in the table below. A non-targeting RNP carrying a control sgRNA ("L2" sgRNA) was used as a control.

| Target Gene Symbol | Expected Deletion Size | sgRNA1 template sequence | PAM | sgRNA2 template sequence | PAM | PCR primar Fwd | PCR primar Rev | PCR fragment | KO Fragment size | Deletion site | Deletion type |
|---|---|---|---|---|---|---|---|---|---|---|---|
| Bcat1 | 214 | GGA TCC TAA TAC GAC TCA CTA TAG CAC GGA TCA TAT GCT GAC GGG TTT TAG AGC TAG AA | TGG | GGA TCC TAA TAC GAC TCA CTA TAG GGC TTT ATC ATA TTT ACC AGG TTT TAG AGC TAG AA | AGG | CAC GGA TCA TAT GCT GAC GG | GGC TTT ATC ATA TTT ACC AG | 550 | 336 | 3-i | Exon-intron pair |

Electroporation was performed with a Neon transfection device (Invitrogen): 100,000 cells were re-suspended in 20 µL buffer R, 10 µL of pre-assembled RNP mix was added and the solution was mixed by careful pipetting. Electroporation was performed in 10 µL Neon-tips using the following parameters (two rounds of electroporation, cells were pooled afterwards): - Voltage: 1'325 V - Pulse width: 10 mS - Pulse number: 3 Cells were expanded and single-cell clones were sorted into 96-well U-bottom plates 7 days after electroporation. Multiple clones were picked and screened for successful knockout (fragment deletion), which was indicated by the presence of a single PCR fragment at ~336 bp. Selected clones were further validated by Sanger sequencing, showing homozygous deletion of the expected fragments in clones of interest. Clones of interest and control clones were expanded, frozen in Bambanker cryoprotection medium (Nalgene) and stored at -80°C.

**Animal Maintenance**. Animals were housed in filter-topped and autoclaved, solid floor polycarbonate cages supplied with autoclaved bedding. One day prior to treatment initiation, the mice were ear-tagged, weighed, shaved, and sorted. The general appearance (thin, fat, dehydrated, etc.), attitude (anesthetized, moribund, cataleptic, depressed, hyperactive, aggressive), movement (ataxia, convulsions, pruritus, tremor, fasciculation, and paralysis), skin/hair (rough hair coat), eyes (discharge and swelling), nose (discharges) and respiration (dyspnea, gasping, etc) were monitored daily throughout the test and recorded, if abnormal. Signs of distress, weight loss more than 20% and tumor volumes ≥2000 $mm^3$ were criteria for euthanasia.

**Bioinformatics**.

*Gene Set Enrichment Analysis* (GSEA v5.2) was utilized to further assess whether specific biological pathways or signatures were significantly enriched between different treatment groups. GSEA determines whether an *a priori* defined gene set shows statistically significant cumulative changes in gene expression between phenotypic subgroups. In brief, all differentially expressed genes were ranked according to the fold change of expression between two groups. Next, an enrichment score was calculated for a given gene set based on its position in the ranked differential list. One thousand random permutations of the phenotypic subgroups were used to establish a null distribution of enrichment score against which a normalized enrichment score (NES) and false discovery rate (FDR)-corrected *q* values were calculated. All gene sets with an FDR less than 0.25 were considered as statistically significant.

*Pathway Enrichment Analysis*. To determine if specific biological pathways were enriched, DEGs (p<0.05 and fold change≥1.5) were utilized in conjunction with the pathfindR package. KEGG (http://www.kegg.jp) is a database of genes and genomes with assigned functional meaning at the molecular and higher level (S3). Molecular level functions are found in the KEGG Orthology database, whereas higher level functions are stored as networks of interactions and reactions. Reactome (https://reactome.org) is an open-source and peer-reviewed database that is manually curated (S4). Pathways analysis was performed using both KEGG and Reactome. P-values were calculated according to the following formula:

$$p = 1 - \sum_{i=0}^{m-1} \frac{\binom{M}{i}\binom{N-M}{n-i}}{\binom{N}{n}},$$

where N is the number of all genes with KEGG or Reactome annotation, n is the number of target gene candidates in N, M is the number of all genes annotated to a certain pathway, and m is the number of target gene candidates in M.

*Cytoscape*. For network visualization of DEGs that were either positively or negatively affected by treatment, we utilized using Cytoscape (v3.8.2) (S5). We applied a minimum adjacency threshold of 0.03 for including edges between genes, and we removed genes that were not connected to the main module network. We clustered proteins in the resulting networks using MCODE (S6). Within a PPI network, MCODE clusters proteins that are part of the same pathways or part of the same protein complexes.

*WGCNA*. The WCGNA method was used to construct modules of coexpressed genes that correlated with genomic phenotype of the cells following ERG245 treatment and to identify hub gene candidates within those modules. This was executed using the WGCNA R package per authors' instructions for automatic network construction and module detection, which typically involves creating a cluster tree, assigning branches as modules, merging close modules, and determining module eigengene (S7). Briefly, the expression of genes belonging to the RNAseq dataset (QD356), obtained after activation of $CD8^+$ T cells in the absence and presence of ERG245 for 24h, was totaled and genes with sum FPKM<60 were excluded from further analysis. *Hierarchical Clustering* of the dataset detected two outliers (EEF1A1, B2M), which were manually removed (Supplementary Fig. S2C). We assumed signed correlation networks and computed bi-weight mid-correlations. Using the *pickSoftThreshold* function and a threshold of 0.9 for the scale-free topology index, we set the soft thresholding power at p=18 for determining gene adjacency (Supplementary Fig. S2D). The function *blockwisemodules* was then used to obtain the modules and the corresponding eigengene values. Cluster dendrograms are shown in Supplementary Fig. S3E. In that figure, colors indicate modules after merging those in close proximity. The least square regression method was then used to identify modules with genes that are differentially expressed after treatment (Supplementary Fig. S3F). Those modules are shown in Supplementary Table S2. The function of each module of Table S2 was then characterized using Gene Ontology (GO) Enrichment Analysis (http://www.geneontology.org). Biological processes exhibiting a combination of high Fold Change (FC) with treatment and low False Discovery Rate (FDR) are listed in Supplementary Table S3. To identify gene hubs within modules of interest, we determined module membership kME values for all genes and all modules. The module membership KME of a gene in a specific module is defined as the Pearson correlation coefficient between the gene and the module eigengene. As hub gene candidates, we selected genes with a high degree of differential expression after treatment (determined by a combination of a high FC value and low FDR value) that also possessed a high KME value within a certain module (Supplementary Table S4). We then examined the known functions of the candidate genes in Table S4 in light of the biological processes linked to the modules in Table S3 and highlighted genes of interest (Table S4).

**Table S1**

| 1 mg/kg, iv | | | | | | | |
|---|---|---|---|---|---|---|---|
| $t_{1/2}$ | $C_0$ | $AUC_{last}$ | $AUC_{inf}$ | $AUC_{Extr}$ | MRT | Vss | CL |
| *(h)* | *(ng/mL)* | *(h*ng/mL)* | *(h*ng/mL)* | *(%)* | *(h)* | *(L/kg)* | *(mL/min/Kg)* |
| 5.23 | 18099 | 48819 | 50501 | 3.33 | 5.96 | 0.12 | 0.33 |
| **10 mg/kg, po** | | | | | | | |
| $t_{max}$ | $C_{max}$ | $AUC_{last}$ | $AUC_{inf}$ | AUC/D | $AUC_{Extr}$ | MRT | F |
| *(h)* | *(ng/mL)* | *(h*ng/mL)* | *(h*ng/mL)* | *(h*kg*ng/mL/mg)* | *(%)* | *(h)* | *(%)* |
| 0.17 | 45587 | 345995 | 356149 | 35615 | 2.85 | 6.32 | 71 |

$t_{1/2}$: half-life
$C_0$: initial serum concentration after iv injection
AUC: area under the curve
MRT: mean residence time
$V_{ss}$: steady-state volume of distribution
$t_{max}$: time to Cmax
$C_{max}$: maximal serum concentration after oral administration
F: bioavailability

**Table S2**

| Module | logFC | P-Value | Number of Genes/Module |
|---|---|---|---|
| ME3 | -0.77 | 0.029 | 830 |
| ME1 | -0.74 | 0.038 | 2069 |
| ME2 | 0.72 | 0.044 | 1211 |
| ME8 | -0.71 | 0.045 | 278 |

**Table S3**

| **GO biological process complete** | **Fold Enrichment** | **FDR** |
|---|---|---|
| **ME3** | | |
| positive regulation of chromosome separation (GO:1905820) | 9.34 | 5.82E-03 |
| secondary alcohol biosynthetic process (GO:1902653) | 6.17 | 6.93E-03 |
| **cholesterol biosynthetic process** (GO:0006695) | 6.17 | 6.87E-03 |
| **oxidative phosphorylation** (GO:0006119) | 4.49 | 4.76E-05 |
| mitochondrial ATP synthesis coupled electron transport (GO:0042775) | 4.48 | 1.15E-03 |
| ATP synthesis coupled electron transport (GO:0042773) | 4.48 | 1.14E-03 |
| respiratory electron transport chain (GO:0022904) | 3.86 | 2.35E-03 |
| **ME1** | | |
| DNA strand elongation involved in DNA replication (GO:0006271) | 10.23 | 7.49E-07 |
| nuclear DNA replication (GO:0033260) | 7.11 | 6.44E-06 |
| maturation of LSU-rRNA (GO:0000470) | 6.94 | 6.59E-07 |
| DNA replication initiation (GO:0006270) | 6.82 | 1.83E-06 |
| cell cycle DNA replication (GO:0044786) | 6.82 | 9.34E-06 |
| spliceosomal snRNP assembly (GO:0000387) | 5.80 | 1.06E-06 |
| maturation of 5.8S rRNA (GO:0000460) | 5.55 | 7.36E-06 |
| DNA-dependent DNA replication (GO:0006261) | 5.03 | 9.27E-18 |
| ribosomal large subunit biogenesis (GO:0042273) | 4.97 | 6.33E-10 |
| **rRNA processing** (GO:0006364) | 4.81 | 3.03E-29 |
| **DNA replication** (GO:0006260) | 4.69 | 2.82E-24 |
| **ribosome biogenesis** (GO:0042254) | 4.64 | 7.17E-37 |
| **rRNA metabolic process** (GO:0016072) | 4.62 | 1.92E-28 |
| ribosomal small subunit biogenesis (GO:0042274) | 4.62 | 1.01E-08 |
| **ribonucleoprotein complex biogenesis** (GO:0022613) | 4.34 | 1.83E-47 |
| RNA splicing, via transesterification reactions (GO:0000375) | 4.08 | 8.42E-23 |
| RNA splicing, via transesterification reactions with bulged adenosine as nucleophile (GO:0000377) | 4.06 | 3.25E-22 |
| mRNA splicing, via spliceosome (GO:0000398) | 4.06 | 3.32E-22 |
| ribonucleoprotein complex localization (GO:0071166) | 3.87 | 3.93E-06 |
| ncRNA processing (GO:0034470) | 3.80 | 2.48E-32 |
| **ME2** | | |
| vesicle fusion with Golgi apparatus (GO:0048280) | 10.49 | 9.94E-03 |
| UV-damage excision repair (GO:0070914) | 9.41 | 4.95E-03 |
| intrinsic apoptotic signaling pathway in response to endoplasmic reticulum stress (GO:0070059) | 5.30 | 6.63E-03 |
| nucleotide-excision repair (GO:0006289) | 5.09 | 1.27E-04 |
| regulation of transcription from RNA polymerase II promoter in response to stress (GO:0043618) | 4.64 | 2.14E-03 |
| **regulation of response to endoplasmic reticulum stress** (GO:1905897) | 4.16 | 9.39E-05 |

**Table S3 (cont.)**

| GO biological process complete | Fold Enrichment | FDR |
|---|---|---|
| cytoplasmic translation (GO:0002181) | 3.30 | 4.11E-04 |
| positive regulation of cellular protein catabolic process (GO:1903364) | 3.25 | 3.66E-05 |
| regulation of cold-induced thermogenesis (GO:0120161) | 3.16 | 2.05E-04 |
| positive regulation of proteolysis involved in cellular protein catabolic process (GO:1903052) | 3.13 | 5.19E-04 |
| **ME8** | | |
| NADH regeneration (GO:0006735) | 37.18 | 5.25E-04 |
| glucose catabolic process to pyruvate (GO:0061718) | 37.18 | 5.09E-04 |
| canonical glycolysis (GO:0061621) | 37.18 | 4.94E-04 |
| glycolytic process through glucose-6-phosphate (GO:0061620) | 30.98 | 8.27E-04 |
| glycolytic process through fructose-6-phosphate (GO:0061615) | 30.98 | 8.07E-04 |
| **glycolytic process** (GO:0006096) | 26.02 | 6.01E-11 |
| ATP generation from ADP (GO:0006757) | 25.39 | 5.31E-11 |
| ADP metabolic process (GO:0046031) | 22.76 | 6.69E-11 |
| nucleoside diphosphate phosphorylation (GO:0006165) | 18.90 | 1.70E-10 |
| nucleotide phosphorylation (GO:0046939) | 18.28 | 2.07E-10 |
| purine ribonucleoside diphosphate metabolic process (GO:0009179) | 17.99 | 2.11E-10 |
| purine nucleoside diphosphate metabolic process (GO:0009135) | 17.99 | 1.81E-10 |
| ribonucleoside diphosphate metabolic process (GO:0009185) | 17.16 | 2.88E-10 |
| pyruvate metabolic process (GO:0006090) | 17.06 | 1.62E-09 |
| carbohydrate catabolic process (GO:0016052) | 10.72 | 7.54E-08 |
| hexose metabolic process (GO:0019318) | 8.26 | 4.08E-07 |
| ATP metabolic process (GO:0046034) | 8.10 | 5.45E-10 |
| monosaccharide metabolic process (GO:0005996) | 7.66 | 3.81E-07 |
| Golgi vesicle transport (GO:0048193) | 4.94 | 2.07E-05 |
| purine ribonucleotide metabolic process (GO:0009150) | 4.13 | 6.68E-04 |
| ribose phosphate metabolic process (GO:0019693) | 3.99 | 5.77E-04 |
| nucleobase-containing small molecule metabolic process (GO:0055086) | 3.62 | 7.59E-05 |
| organophosphate metabolic process (GO:0019637) | 3.39 | 1.24E-07 |
| monocarboxylic acid metabolic process (GO:0032787) | 3.33 | 6.40E-04 |

**Table S4**

| Name | kME | FC | FDR | Name | kME | FC | FDR | Name | kME | FC | FDR |
|---|---|---|---|---|---|---|---|---|---|---|---|
| **ME3** | | | | **ME1** | | | | **ME2** | | | |
| IDI1 | 0.997 | -1.76 | 4.88E-04 | CCT8 | 1.000 | -1.68 | 7.53E-04 | JMY | 0.998 | 2.12 | 4.92E-05 |
| DDX1 | 0.994 | -1.55 | 5.72E-03 | RANBP1 | 0.997 | -1.83 | 1.58E-04 | ARL4C | 0.997 | 1.83 | 7.68E-04 |
| YWHAB | 0.991 | -1.61 | 2.81E-03 | TOMM5 | 0.997 | -1.72 | 5.21E-04 | ARRDC3 | 0.992 | 2.98 | 9.09E-08 |
| BZW1 | 0.988 | -1.57 | 1.75E-03 | MAGOHB | 0.997 | -1.85 | 1.02E-03 | NXF1 | 0.992 | 1.87 | 8.70E-05 |
| MATK | 0.987 | -2.43 | 4.39E-06 | HSPE1 | 0.996 | -2.10 | 7.59E-06 | ***ISCU*** | *0.992* | *1.96* | *1.02E-04* |
| ACTG1 | 0.983 | -2.15 | 2.17E-06 | TUBA1B | 0.996 | -2.55 | 6.21E-06 | DDIT3 | 0.952 | 3.88 | 8.38E-08 |
| MRPS27 | 0.983 | -1.55 | 6.01E-03 | EIF5A | 0.995 | -1.92 | 2.25E-05 | WSB1 | 0.962 | 2.41 | 3.91E-05 |
| SCFD2 | 0.982 | -2.10 | 3.82E-04 | ACOT7 | 0.995 | -2.24 | 3.21E-05 | **ME8** | | | |
| EEF2K | 0.982 | -1.57 | 4.70E-03 | CYCS | 0.995 | -1.94 | 8.93E-05 | VDR | 0.991 | -2.33 | 2.04E-05 |
| MSMO1 | 0.980 | -1.78 | 9.13E-05 | VDAC1 | 0.995 | -1.98 | 1.66E-05 | AK4 | 0.968 | -6.35 | 1.21E-08 |
| KCNK5 | 0.980 | -1.75 | 2.74E-03 | NME1 | 0.994 | -2.15 | 7.59E-06 | EGR3 | 0.960 | -4.75 | 1.41E-10 |
| PPA1 | 0.980 | -1.67 | 1.77E-03 | ***CISD1*** | *0.985* | *-2.73* | *1.01E-05* | HILPDA | 0.955 | -3.68 | 4.13E-05 |
| MLKL | 0.979 | -1.70 | 6.46E-03 | MCM2 | 0.981 | -4.52 | 1.52E-08 | **LDHA** | *0.983* | *-3.26* | *1.51E-11* |
| CALR | 0.978 | -1.52 | 6.46E-03 | HPDL | 0.980 | -4.00 | 9.92E-06 | TMC8 | 0.967 | -2.36 | 4.90E-08 |
| GTF3C6 | 0.978 | -1.51 | 8.33E-03 | MCM4 | 0.977 | -6.94 | 4.51E-13 | PFKFB3 | 0.956 | -2.24 | 4.70E-04 |
| MYCBP | 0.976 | -1.54 | 5.17E-03 | CHEK1 | 0.976 | -7.96 | 1.10E-12 | GBE1 | 0.954 | -2.15 | 5.12E-04 |
| ATP5G3 | 0.976 | -1.73 | 1.30E-03 | FEN1 | 0.971 | -3.97 | 6.24E-11 | PGAM1 | 0.962 | -2.09 | 2.52E-05 |
| ***BCAT1*** | *0.974* | *-2.23* | *1.14E-04* | CDC6 | 0.971 | -8.68 | 5.32E-14 | PGK1 | 0.985 | -1.95 | 1.14E-04 |
| LTA | 0.973 | -2.89 | 1.12E-06 | GINS2 | 0.970 | -9.93 | 5.15E-06 | ALDOA | 0.958 | -1.92 | 2.43E-06 |
| NUP210 | 0.971 | -1.55 | 5.22E-03 | ZNF367 | 0.970 | -6.39 | 1.88E-10 | TPI1 | 0.974 | -1.91 | 4.84E-04 |
| BRCC3 | 0.971 | -1.66 | 2.19E-03 | CDT1 | 0.969 | -9.63 | 1.64E-06 | PFKP | 0.977 | -1.89 | 1.15E-04 |
| PTPN7 | 0.968 | -1.65 | 9.08E-03 | GINS1 | 0.969 | -6.55 | 1.01E-08 | ENO1 | 0.969 | -1.84 | 1.44E-05 |
| TFAM | 0.966 | -1.58 | 3.58E-03 | ORC1 | 0.968 | -12.08 | 3.28E-15 | MTFP1 | 0.956 | -1.65 | 4.25E-03 |
| ACAT2 | 0.965 | -2.61 | 2.82E-07 | PLK4 | 0.966 | -5.20 | 1.20E-06 | ICAM1 | 0.977 | -1.56 | 8.31E-03 |
| ARL5A | 0.965 | -1.51 | 6.46E-03 | E2F1 | 0.965 | -4.36 | 5.03E-09 | | | | |
| C17orf96 | 0.961 | -1.91 | 7.60E-04 | TYMS | 0.960 | -5.40 | 2.18E-10 | | | | |
| BRK1 | 0.960 | -1.65 | 1.86E-03 | CLSPN | 0.959 | -6.60 | 7.55E-12 | | | | |
| KCNN4 | 0.959 | -1.77 | 6.21E-03 | HELLS | 0.958 | -4.36 | 1.15E-08 | | | | |
| EGR1 | 0.957 | -3.78 | 8.17E-11 | CHAF1A | 0.957 | -4.73 | 1.21E-08 | | | | |
| EIF3CL | 0.956 | -1.62 | 1.63E-03 | DTL | 0.954 | -15.68 | 2.26E-07 | | | | |

#### Table S5

| DEGs that Reverse Direction of Expression | | | | | DEGs that Maintain Direction of Expression | | | | |
|---|---|---|---|---|---|---|---|---|---|
| Gene | 24h | | 72h | | Gene | 24h | | 72h | |
| | FC | FDR | FC | FDR | | FC | FDR | FC | FDR |
| LTB | -2.56 | 4.15E-04 | 1.41 | 8.08E-05 | **MKI67** | *-9.21* | *3.57E-10* | *-1.93* | *2.94E-05* |
| ALDOC | -2.54 | 4.81E-04 | 1.46 | 2.47E-03 | ZNF367 | -6.39 | 1.88E-10 | -1.42 | 2.99E-03 |
| BNIP3 | -2.30 | 5.25E-03 | 1.39 | 1.81E-03 | ATAD5 | -3.23 | 1.67E-05 | -1.82 | 5.69E-07 |
| GZMB | -2.26 | 4.46E-03 | 1.38 | 7.09E-04 | WEE1 | -3.04 | 6.85E-08 | -1.36 | 4.37E-03 |
| NME1 | -2.15 | 7.59E-06 | 1.38 | 9.33E-04 | KNTC1 | -2.57 | 1.08E-05 | -1.40 | 1.36E-04 |
| IFITM3 | -2.11 | 6.38E-03 | 1.37 | 3.21E-03 | **BCAT1** | *-2.23* | *1.14E-04* | *-1.40* | *3.08E-03* |
| ATP5G1 | -2.11 | 2.78E-05 | 1.38 | 4.69E-03 | MMS22L | -2.00 | 3.72E-04 | -1.40 | 2.84E-03 |
| NPM3 | -2.11 | 2.78E-05 | 1.45 | 9.59E-03 | IPO11 | -1.90 | 3.21E-05 | -1.35 | 3.72E-03 |
| TESC | -2.00 | 4.94E-03 | 1.40 | 5.07E-03 | PM20D2 | -1.85 | 2.50E-04 | -1.44 | 3.39E-04 |
| ALDOA | -1.92 | 2.43E-06 | 1.39 | 9.63E-03 | TOPBP1 | -1.84 | 2.25E-04 | -1.49 | 2.78E-06 |
| TXNDC17 | -1.89 | 3.99E-03 | 1.36 | 2.27E-03 | DHX33 | -1.74 | 2.01E-03 | -1.42 | 1.92E-04 |
| MRPL12 | -1.88 | 6.37E-05 | 1.41 | 1.42E-03 | UTP20 | -1.64 | 3.36E-03 | -1.35 | 3.01E-03 |
| GLA | -1.84 | 4.70E-03 | 1.37 | 2.80E-03 | HCFC1 | -1.60 | 7.23E-03 | -1.50 | 7.04E-08 |
| FAM162A | -1.81 | 6.02E-04 | 1.55 | 5.60E-05 | EPT1 | -1.58 | 4.46E-03 | -1.56 | 2.50E-06 |
| PRDX1 | -1.71 | 2.74E-04 | 1.37 | 5.06E-04 | PRKDC | -1.55 | 8.71E-03 | -1.54 | 5.03E-06 |
| C1QBP | -1.66 | 5.78E-03 | 1.36 | 2.01E-03 | HLA-F | 1.61 | 9.19E-03 | 1.35 | 6.10E-03 |
| MTFP1 | -1.65 | 4.25E-03 | 1.43 | 1.00E-03 | GPX4 | 1.61 | 5.88E-03 | 1.47 | 1.45E-04 |
| MIF | -1.63 | 8.55E-03 | 1.48 | 8.27E-05 | DHRS3 | 1.71 | 1.02E-03 | 1.43 | 5.13E-04 |
| HSD17B10 | -1.59 | 4.09E-03 | 1.38 | 1.12E-03 | ZFAS1 | 1.91 | 1.52E-03 | 1.45 | 6.83E-04 |
| NDUFS5 | -1.58 | 3.56E-03 | 1.39 | 6.12E-05 | IGFLR1 | 1.93 | 1.49E-04 | 1.39 | 6.45E-03 |
| RUVBL2 | -1.52 | 5.22E-03 | 1.37 | 2.36E-04 | PPDPF | 1.93 | 6.83E-06 | 1.35 | 3.23E-04 |
| CRY1 | 1.51 | 5.04E-03 | -1.38 | 1.65E-03 | GAS5 | 2.22 | 9.05E-04 | 1.42 | 3.81E-04 |
| NBPF14 | 1.53 | 3.82E-03 | -2.07 | 1.92E-03 | | | | | |
| HECA | 1.53 | 6.18E-03 | -1.45 | 1.15E-03 | | | | | |
| ZCCHC11 | 1.54 | 5.45E-03 | -1.35 | 2.00E-03 | | | | | |
| KIAA1109 | 1.54 | 5.88E-03 | -2.07 | 2.08E-12 | | | | | |
| MORC3 | 1.54 | 8.02E-03 | -1.43 | 5.60E-04 | | | | | |
| TEP1 | 1.54 | 9.71E-03 | -1.37 | 3.21E-03 | | | | | |
| KCNA3 | 1.58 | 5.64E-03 | -1.60 | 1.63E-04 | | | | | |
| OGT | 1.58 | 1.47E-03 | -1.45 | 3.52E-05 | | | | | |
| GK5 | 1.60 | 3.56E-03 | -1.58 | 5.10E-04 | | | | | |
| NBPF12 | 1.61 | 3.20E-03 | -2.73 | 2.55E-04 | | | | | |
| CHD2 | 1.61 | 2.86E-03 | -1.44 | 1.34E-04 | | | | | |
| RASGRF2 | 1.73 | 3.57E-03 | -1.46 | 5.02E-03 | | | | | |
| SLFN5 | 1.74 | 1.45E-04 | -1.45 | 5.08E-06 | | | | | |
| CREBBP | 1.74 | 2.39E-03 | -1.49 | 1.17E-06 | | | | | |

**Table S5 (cont.)**

| DEGs that Reverse Direction of Expression | | | | | DEGs that Maintain Direction of Expression | | | | |
|---|---|---|---|---|---|---|---|---|---|
| Gene | 24H | | 72H | | Gene | 24H | | 72H | |
| | FC | FDR | FC | FDR | | FC | FDR | FC | FDR |
| ZMAT3 | 1.77 | 5.07E-03 | -1.49 | 1.46E-04 | | | | | |
| KIAA0355 | 1.82 | 3.10E-04 | -1.39 | 9.72E-03 | | | | | |
| GOLGB1 | 1.88 | 1.59E-04 | -1.49 | 9.86E-05 | | | | | |
| DGKH | 1.88 | 9.42E-03 | -1.76 | 5.47E-04 | | | | | |
| AHR | 1.90 | 7.60E-04 | -1.43 | 2.28E-05 | | | | | |
| ATM | 2.02 | 4.05E-05 | -1.62 | 3.83E-07 | | | | | |
| NR1D2 | 2.09 | 1.54E-03 | -1.52 | 2.22E-03 | | | | | |
| FNIP1 | 2.11 | 2.49E-05 | -1.59 | 4.83E-07 | | | | | |
| JMY | 2.12 | 4.92E-05 | -1.51 | 2.14E-03 | | | | | |
| GPCPD1 | 2.13 | 3.75E-05 | -1.60 | 2.44E-04 | | | | | |
| TXK | 2.15 | 2.57E-03 | -1.51 | 1.24E-04 | | | | | |
| SYNE2 | 2.41 | 2.23E-06 | -1.84 | 1.56E-07 | | | | | |
| ARRDC3 | 2.98 | 9.09E-08 | -1.55 | 1.47E-03 | | | | | |

**Table Legends**

**Table S1**. Pharmacokinetic properties of ERG245.

**Table S2**. Gene modules identified using the WGCNA algorithm as significantly impacted by treatment with ERG245.

**Table S3**. GO analyses of gene modules shown in Table S2. Biological processes exhibiting a combination of high Fold Enrichment values with treatment and low FDR values are listed.

**Table S4**. Lists of potential hub genes present in the MEs of Table S2. Identification is based on a combination of a high module membership kME value and a high degree of differential expression after treatment (as determined by high |FC| and low FDR values).

**Table S5**. Lists of common genes, differentially expressed in the RNAseq datasets of $CD8^+$ T cells at 24h (see Fig. 4) and 72h (See Fig. 5).

#### Figure Legends

**Figure S1**. (A) BCAT1 protein expression in naïve, control, and ERG245-treated $hCD8^+$ T cells. Two individual donors. (B) qPCR data analyzing Bcat1 and Bcat2 gene expression in naïve and control $hCD8^+$ T cells. (C) Enzymatic reaction catalyzed by BCAT1. (D) Relative levels of phosphorylated mTOR (p-mTOR) and phosphorylated ribosomal S6 (p-rS6) proteins present in naïve (N), control (C), and ERG245-treated (E) $hCD8^+$ T cells (two individual donors per test; D# indicates specific donors). (E) Levels of c-Myc found in naïve, control, and ERG245-treated $hCD8^+$ T cells (two individual donors). (F) The extracellular levels of select metabolites found in the CM of $hCD8^+$ after exposure to ERG245 for 24h (left) and in the CM of $mCD8^+$ T cells isolated from wt and bcat1 KO animals and cultured for 24h (right). The experiments included 4 individual human donors and 5 age-matched pairs of animals. (G) Ratio of (m+3)/(m+0) Lac found in CL of $hCD8^+$ T cultured in media containing of U-$^{13}$C-Glucose for 24H. Three individual donors. (H) Comparison of the (m+3)/(m+0) Lac ratios found in the CL and CM of $hCD8^+$ T cells after 24h treatment with ERG245 (24h) and after withdrawal of BCAT1i and additional incubation in complete media containing IL-2 for another 24h (48h) (3 individual donors). The experimental design is also shown. (I) Basal respiration, maximal respiration, space respiratory capacity, rate of ATP produced through glycolysis, and rate of ATP produced through oxidative phosphorylation in $hCD8^+$ cells exposed to ERG245 for 24h. (F) Basal respiration, maximal respiration, space respiratory capacity, rate of ATP produced through glycolysis, and rate of ATP produced through oxidative phosphorylation in $hCD8^+$ cells after brief exposure to ERG245 for 24h, withdrawal of BCAT1 inhibition, and additional culturing for 72h (96h).

**Figure S2**. (A) A graphical representation of RNAseq data (collected at 24h and having |FC|≥1.5 and FDR<0.05) combining hierarchical clustering and heatmap. *Red* indicates high gebe expression and *Green* indicates low gene expression. (B) A dotplot showing the top 15 Reactome pathways found to be significantly altered after ERG245 treatment. (C) Gene clustering in WGCNA analysis for identifying outliers. (D) Determination of gene adjacency. (E) Cluster dendrograms obtained after merging modules in close proximity. Colors identify final modules. (F) An example of using the least square regression method to identify modules with genes that were differentially expressed following treatment with ERG245. (G) Immunoprecipitation experiment of rhBCAT1 inhibited with ERG245 and complexed with rhACO1 in a solution containing $(NH_4)Fe(SO_4)_2$ and cysteine. IP: immunoprecipitation. IB: immunoblotting.

**Figure S3**. (A) A graphical representation of RNAseq data (collected at 72h and having |FC|≥1.5 and FDR<0.05) combining hierarchical clustering and heatmap. *Red* indicates a high gene expression and *Blue* indicates a low gene expression. (B) GSEA analysis of all genes detected in the RNAseq data dataseta at 24h and 72h. Enrichments plots of genes in significantly upregulated or downregulated biological pathways of the Hallmark MSigDB collection with FDR<0.05 are shown. The analysis suggests a genomic phenotype reversal with withdrawal of BCAT1i.

**Figure S4**. (A) Frequency and levels of Bcat1 and Bcat2 expression in various cell type clusters as visualized from sc-RNAseq data obtained from CRC, OvC and BC patients (41). (B,C) Normalized graphs showing levels (B) and cell frequency (C) of Bcat1 expression in various T sub-populations found in the TME of lung cancer patients. The graph illustrates that $T_{ex}$ cells preferentially expressed Bcat1 compared to $T_{eff}$ cells. (D) Violin plots showing cytoxicity and exhaustion markers used for the identification of various T cell subpopulations in the TME.

### Figure S1

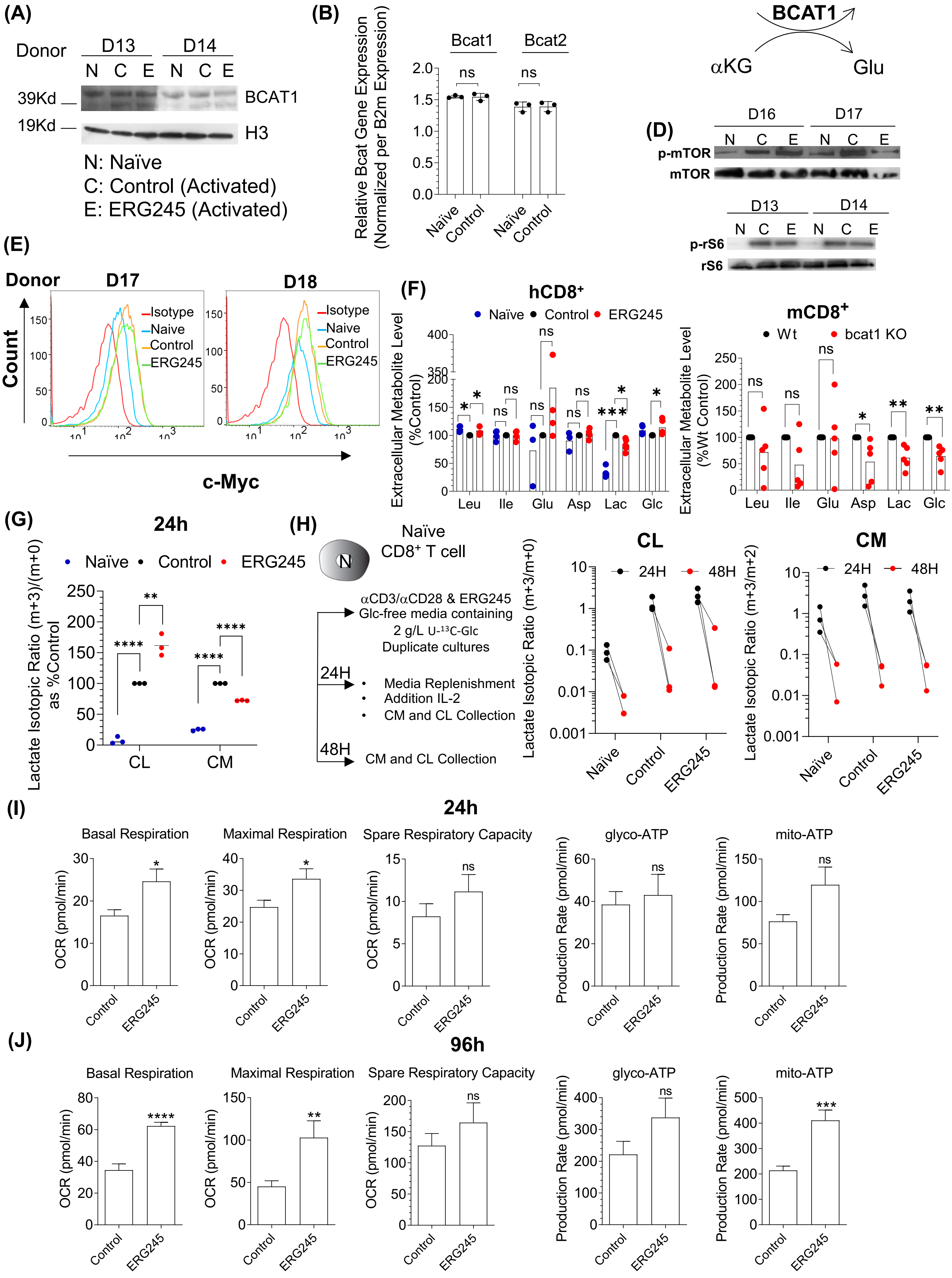
(A)
Donor
D13
D14
N C E N C E
39Kd
19Kd
BCAT1
H3
N: Naïve
C: Control (Activated)
E: ERG245 (Activated)
(B)
Relative Bcat Gene Expression (Normalized per B2m Expression)
Bcat1
Bcat2
ns
Naïve
Control
(C)
Leu
αKIC
BCAT1
αKG
Glu
(D)
D16
D17
p-mTOR
mTOR
D13
D14
p-rS6
rS6
(E)
Donor
D17
D18
Count
c-Myc
Isotype
Naive
Control
ERG245
(F)
hCD8+
Naïve
Control
ERG245
Extracellular Metabolite Level (%Control)
mCD8+
Wt
bcat1 KO
Extracellular Metabolite Level (%Wt Control)
Leu
Ile
Glu
Asp
Lac
Glc
(G)
24h
Lactate Isotopic Ratio (m+3)/(m+0) as %Control
CL
CM
(H)
Naïve CD8+ T cell
αCD3/αCD28 & ERG245
Glc-free media containing
2 g/L U-13C-Glc
Duplicate cultures
24H
Media Replenishment
Addition IL-2
CM and CL Collection
48H
CM and CL Collection
CL
CM
24H
48H
Lactate Isotopic Ratio (m+3/m+0)
Lactate Isotopic Ratio (m+3/m+2)
(I)
24h
Basal Respiration
Maximal Respiration
Spare Respiratory Capacity
glyco-ATP
mito-ATP
OCR (pmol/min)
Production Rate (pmol/min)
Control
ERG245
(J)
96h
Basal Respiration
Maximal Respiration
Spare Respiratory Capacity
glyco-ATP
mito-ATP

### Figure S2

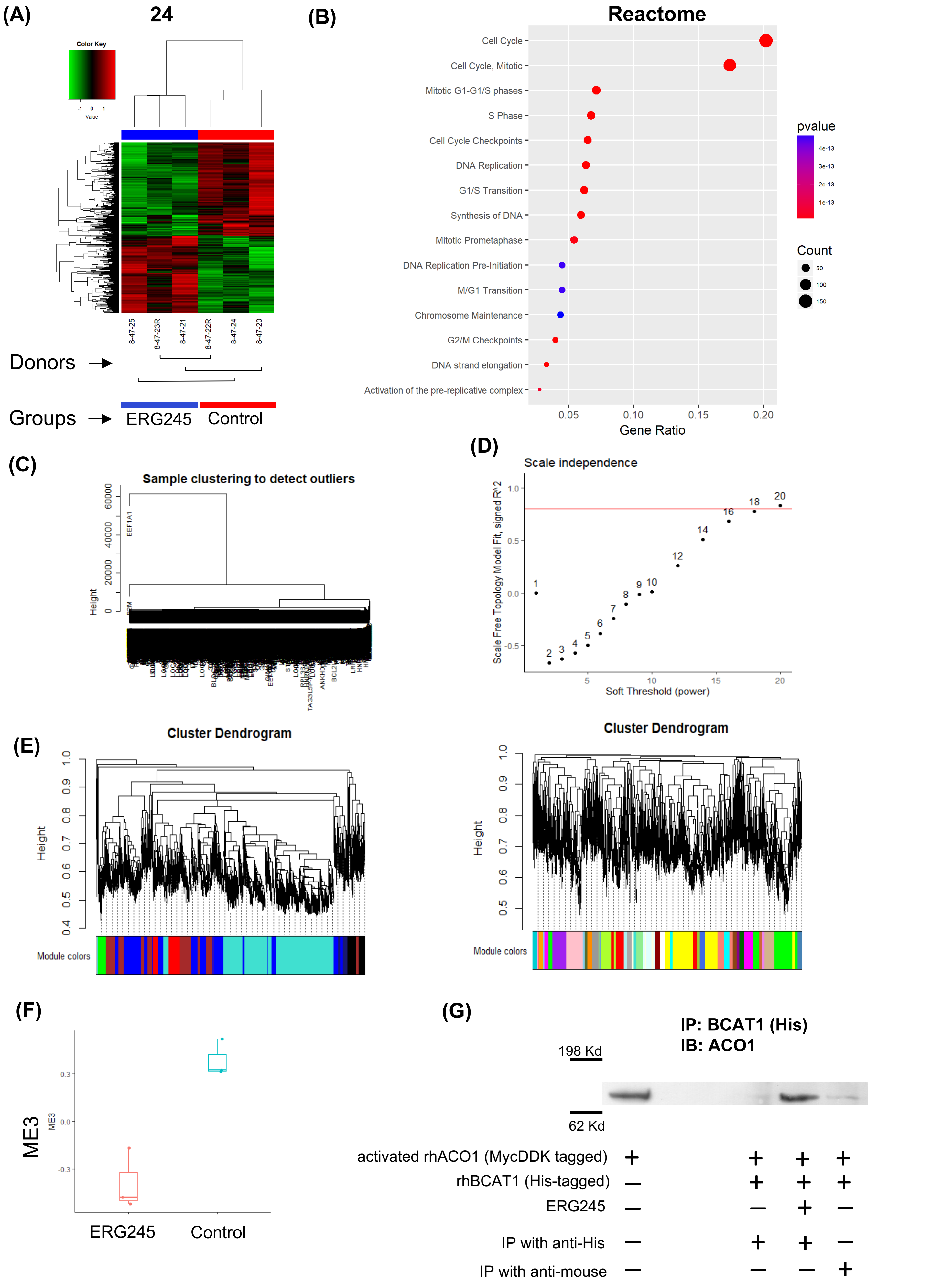
(A)
24
Color Key
Value
8-47-25
8-47-23R
8-47-21
8-47-22R
8-47-24
8-47-20
Donors
Groups
ERG245
Control
(B)
Reactome
Cell Cycle
Cell Cycle, Mitotic
Mitotic G1-G1/S phases
S Phase
Cell Cycle Checkpoints
DNA Replication
G1/S Transition
Synthesis of DNA
Mitotic Prometaphase
DNA Replication Pre-Initiation
M/G1 Transition
Chromosome Maintenance
G2/M Checkpoints
DNA strand elongation
Activation of the pre-replicative complex
Gene Ratio
pvalue
Count
(C)
Sample clustering to detect outliers
Height
EEF1A1
(D)
Scale independence
Scale Free Topology Model Fit, signed R^2
Soft Threshold (power)
(E)
Cluster Dendrogram
Height
Module colors
Cluster Dendrogram
Height
Module colors
(F)
ME3
ERG245
Control
(G)
IP: BCAT1 (His)
IB: ACO1
198 Kd
62 Kd
activated rhACO1 (MycDDK tagged)
rhBCAT1 (His-tagged)
ERG245
IP with anti-His
IP with anti-mouse

### Figure S3

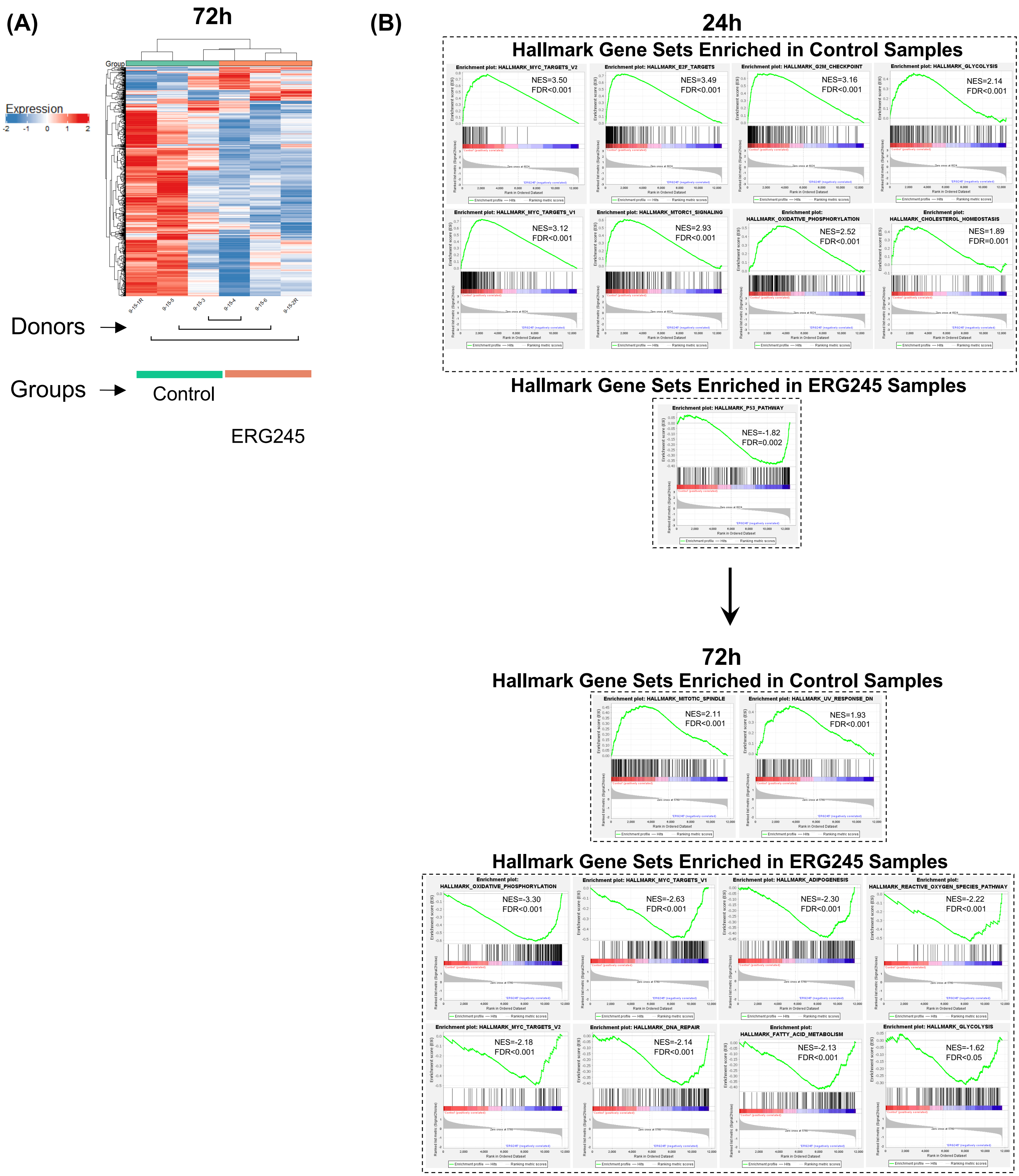

(A)
72h
Group
Expression
-2 -1 0 1 2
9-15-1R
9-15-5
9-15-3
9-15-4
9-15-6
9-15-2R
Donors
Groups
Control
ERG245
(B)
24h
Hallmark Gene Sets Enriched in Control Samples
Enrichment plot: HALLMARK_MYC_TARGETS_V2
NES=3.50
FDR<0.001
Enrichment plot: HALLMARK_E2F_TARGETS
NES=3.49
FDR<0.001
Enrichment plot: HALLMARK_G2M_CHECKPOINT
NES=3.16
FDR<0.001
Enrichment plot: HALLMARK_GLYCOLYSIS
NES=2.14
FDR<0.001
Enrichment plot: HALLMARK_MYC_TARGETS_V1
NES=3.12
FDR<0.001
Enrichment plot: HALLMARK_MTORC1_SIGNALING
NES=2.93
FDR<0.001
Enrichment plot: HALLMARK_OXIDATIVE_PHOSPHORYLATION
NES=2.52
FDR<0.001
Enrichment plot: HALLMARK_CHOLESTEROL_HOMEOSTASIS
NES=1.89
FDR=0.001
Hallmark Gene Sets Enriched in ERG245 Samples
Enrichment plot: HALLMARK_P53_PATHWAY
NES=-1.82
FDR=0.002
72h
Hallmark Gene Sets Enriched in Control Samples
Enrichment plot: HALLMARK_MITOTIC_SPINDLE
NES=2.11
FDR<0.001
Enrichment plot: HALLMARK_UV_RESPONSE_DN
NES=1.93
FDR<0.001
Hallmark Gene Sets Enriched in ERG245 Samples
Enrichment plot: HALLMARK_OXIDATIVE_PHOSPHORYLATION
NES=-3.30
FDR<0.001
Enrichment plot: HALLMARK_MYC_TARGETS_V1
NES=-2.63
FDR<0.001
Enrichment plot: HALLMARK_ADIPOGENESIS
NES=-2.30
FDR<0.001
Enrichment plot: HALLMARK_REACTIVE_OXYGEN_SPECIES_PATHWAY
NES=-2.22
FDR<0.001
Enrichment plot: HALLMARK_MYC_TARGETS_V2
NES=-2.18
FDR<0.001
Enrichment plot: HALLMARK_DNA_REPAIR
NES=-2.14
FDR<0.001
Enrichment plot: HALLMARK_FATTY_ACID_METABOLISM
NES=-2.13
FDR<0.001
Enrichment plot: HALLMARK_GLYCOLYSIS
NES=-1.62
FDR<0.05

### Figure S4

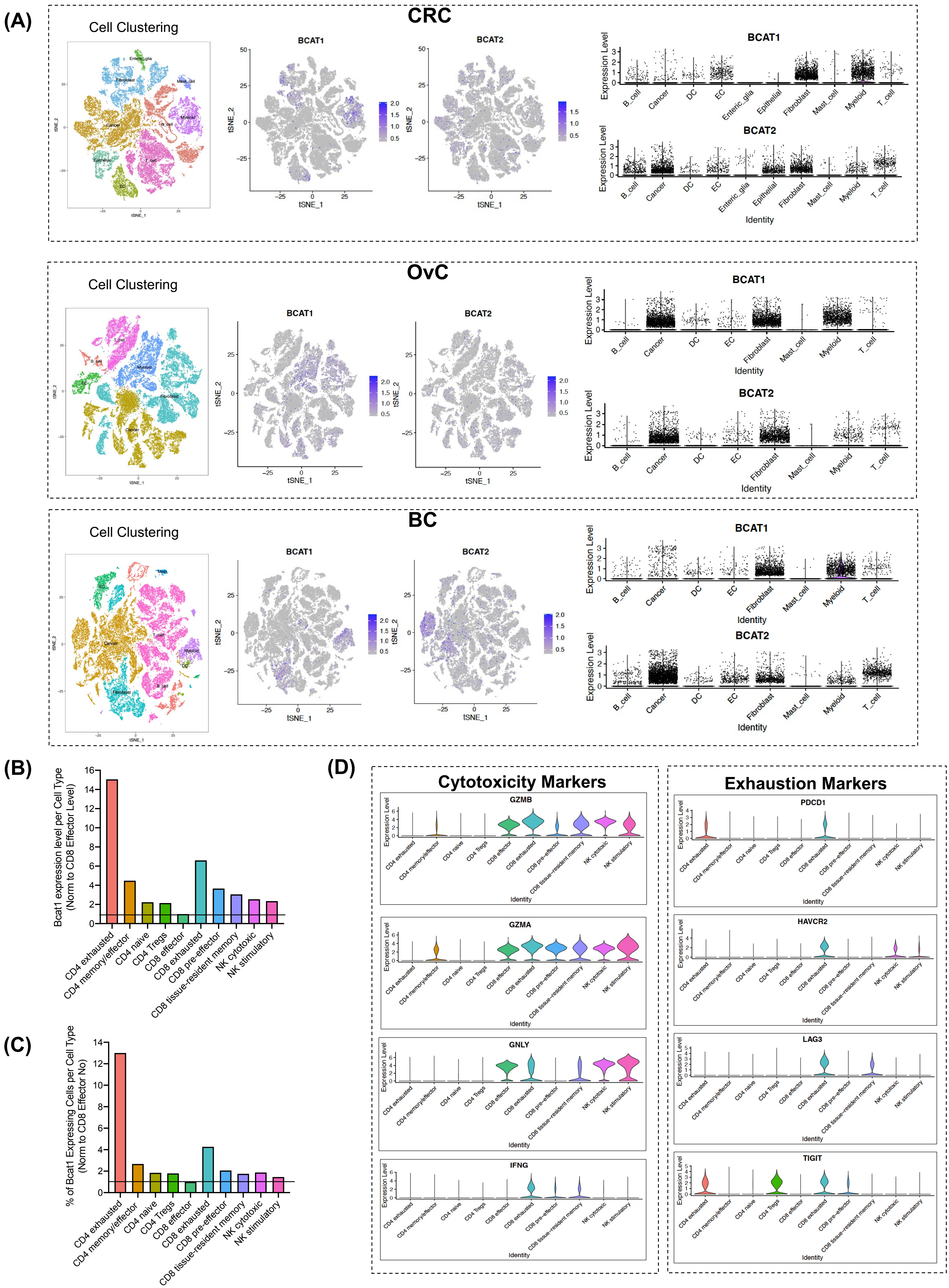

(A)
CRC
Cell Clustering
BCAT1
BCAT2
tSNE_1
tSNE_2
Expression Level
B_cell
Cancer
DC
EC
Enteric_glia
Epithelial
Fibroblast
Mast_cell
Myeloid
T_cell
Identity
OvC
BC
(B)
Bcat1 expression level per Cell Type (Norm to CD8 Effector Level)
(C)
% of Bcat1 Expressing Cells per Cell Type (Norm to CD8 Effector No)
CD4 exhausted
CD4 memory/effector
CD4 naive
CD4 Tregs
CD8 effector
CD8 exhausted
CD8 pre-effector
CD8 tissue-resident memory
NK cytotoxic
NK stimulatory
(D)
Cytotoxicity Markers
GZMB
GZMA
GNLY
IFNG
Exhaustion Markers
PDCD1
HAVCR2
LAG3
TIGIT
